# Generative design of novel bacteriophages with genome language models

**DOI:** 10.1101/2025.09.12.675911

**Authors:** Samuel H. King, Claudia L. Driscoll, David B. Li, Daniel Guo, Aditi T. Merchant, Garyk Brixi, Max E. Wilkinson, Brian L. Hie

**Affiliations:** Arc Institute, Palo Alto, CA, USA; Department of Bioengineering, Stanford University, Stanford, CA, USA; Department of Chemical Engineering, Stanford University, Stanford, CA, USA; Department of Computer Science, Stanford University, Stanford, CA, USA; Department of Genetics, Stanford University, Stanford, CA, USA; Structural Biology Program, Memorial Sloan Kettering Cancer Center, New York, NY, USA; Stanford Data Science, Stanford University, Stanford, CA, USA

## Abstract

Many important biological functions arise not from single genes, but from complex interactions encoded by entire genomes. Genome language models have emerged as a promising strategy for designing biological systems, but their ability to generate functional sequences at the scale of whole genomes has remained untested. Here, we report the first generative design of viable bacteriophage genomes. We leveraged frontier genome language models, Evo 1 and Evo 2, to generate whole-genome sequences with realistic genetic architectures and desirable host tropism, using the lytic phage ΦX174 as our design template. Experimental testing of AI-generated genomes yielded 16 viable phages with substantial evolutionary novelty. Cryo-electron microscopy revealed that one of the generated phages utilizes an evolutionarily distant DNA packaging protein within its capsid. Multiple phages demonstrate higher fitness than ΦX174 in growth competitions and in their lysis kinetics. A cocktail of the generated phages rapidly overcomes ΦX174-resistance in three *E. coli* strains, demonstrating the potential utility of our approach for designing phage therapies against rapidly evolving bacterial pathogens. This work provides a blueprint for the design of diverse synthetic bacteriophages and, more broadly, lays a foundation for the generative design of useful living systems at the genome scale.

## 1. Introduction

Advances in DNA sequencing and synthesis have vastly improved our ability to read and write DNA at the scale of whole genomes (Edgar et al., 2022; Gibson et al., 2008; Hutchison et al., 2016; Richardson et al., 2017; K. Wang et al., 2019). While these technologies could also facilitate whole-genome design, potentially enabling new biotechnologies or therapeutic modalities (Dedrick et al., 2019; Durrant et al., 2024; Mandell et al., 2015; Nyerges et al., 2023), intelligently composing novel genomes still represents a monumental challenge. Genomes contain emergent complexity, wherein interactions involving multiple genes, regulatory elements, recognition sequences, and other features are tightly orchestrated to enable replication and other higher-order functions (**Figure 1A**) (Costanzo et al., 2016; Elena & Lenski, 1997; Jacob & Monod, 1961). Genomes are also highly sensitive to their sequence composition, as even a single mutation can render an entire genome nonviable (Barrell et al., 1976; Hutchison et al., 1999; Sanjuán et al., 2004). Despite efforts to design novel proteins, reprogram genetic codes, and engineer increasingly complex biological circuits (Dauparas et al., 2022; Fredens et al., 2019; Hayes et al., 2025; Hie et al., 2024; Ingraham et al., 2023; Jiang et al., 2024; Srinivasan & Smolke, 2020), there remains no generalizable framework for designing entire genomes.

**Figure 1.**
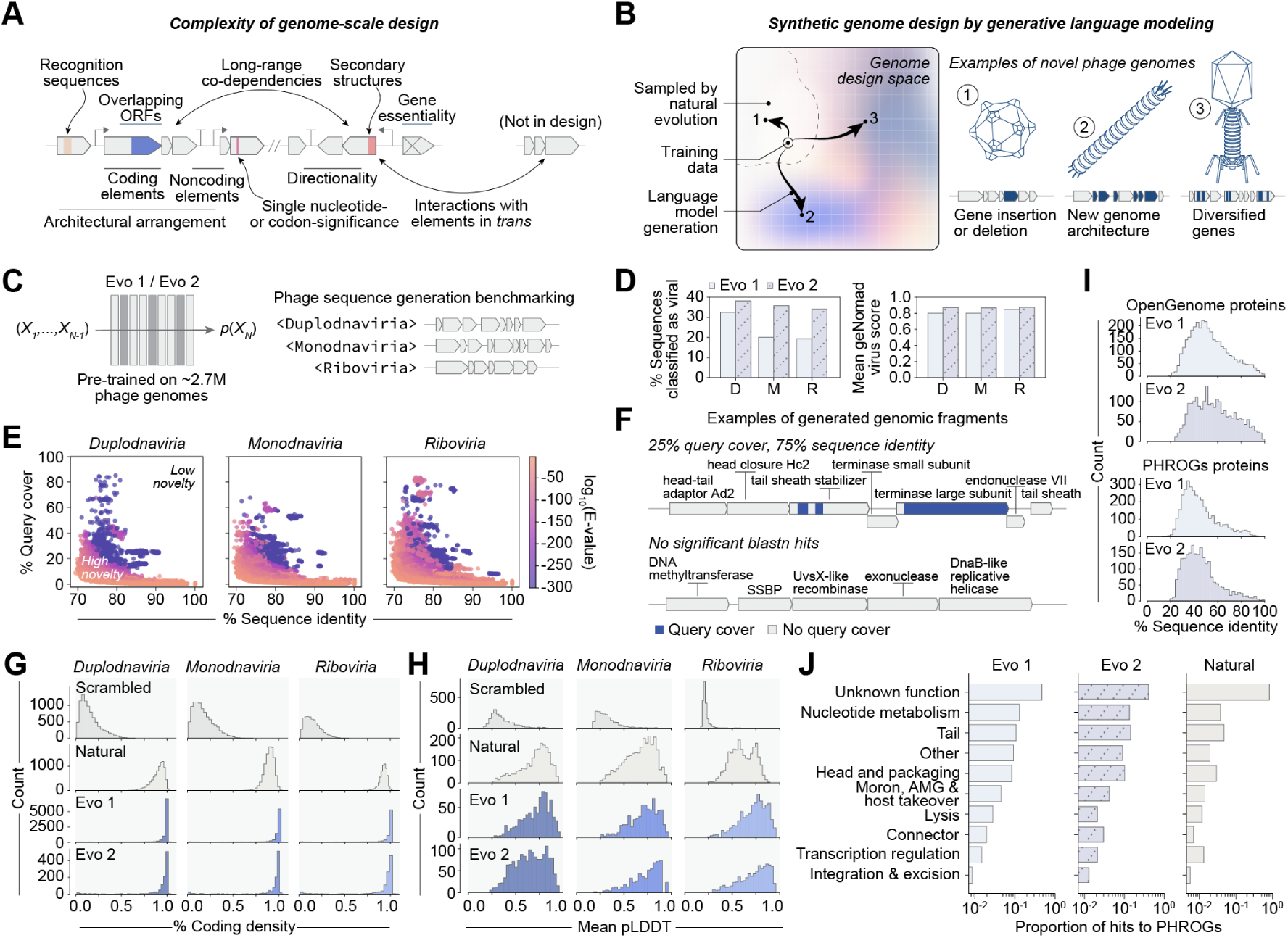
Evo composes realistic bacteriophage genomic sequences. **(A)** Key considerations highlighting the complexity of genome-scale design. **(B)** Generative genome language models have the potential to access novel phage genome design space when trained on a subset of observed natural evolution. **(C)** We benchmarked Evo 1 and Evo 2 on a broad zero-shot prompting task for phage genome design. **(D)** Generated sequences consistently classified as viral by geNomad (left), with high average virus classification scores across prompts (right). D, *Duplodnaviria*; M, *Monodnaviria*; R, *Riboviria*. **(E)** Generated sequences show low query cover and sequence identity against natural sequences in nucleotide BLAST searches, indicating high novelty. **(F)** Generated sequences contain predicted phage-like architectures, with regions that match natural sequences (blue) and novel regions without nucleotide BLAST hits (gray). **(G)** Predicted coding densities of generated sequences are high, similar to natural sequences and unlike scrambled natural sequences. **(H)** ESMFold-predicted protein structures from generated sequences have mean predicted local distance difference test (pLDDT) scores similar to natural proteins, substantially higher than scrambled natural sequence controls. **(I)** Generated proteins align to proteins in the OpenGenome and PHROGs databases with generally low sequence identities, indicating high novelty. **(J)** Functional annotations of generated proteins closely match those of natural phages when queried against the PHROGs database.

Recent advances in artificial intelligence leverage data and compute at scale to enable complex generative tasks (Kaplan et al., 2020; Vaswani et al., 2017). In biology, language models trained on vast genomic sequence datasets learn complex rules that enable the generative design of novel DNA sequences with desired functions (Brixi et al., 2025; Merchant et al., 2024; Nguyen, Poli, Durrant, et al., 2024). For example, we have previously reported the successful design of functional systems such as CRISPR-Cas complexes, transposable elements, and toxin-antitoxin interactions with the genomic language models, Evo 1 and Evo 2 (Merchant et al., 2024; Nguyen, Poli, Durrant, et al., 2024). However, these systems contain only a small number of genetic components, whereas the design of complete genomes would require new methodological advances and much deeper integration of machine learning, computational biology, and experimental biology.

Here, we report the first generative design of complete genomes. We focused on the design of bacteriophages, which have tremendous utility as biotechnological tools and have therapeutic relevance as treatments for bacterial infections (Dedrick et al., 2019; Kilcher & Loessner, 2019; Kim et al., 2025; Pires et al., 2016; Strathdee et al., 2023). As a design template, we selected ΦX174, a lytic *Microviridae* phage with a ∼5.4 kilobase (kb) genome containing 11 genes, at least seven regulatory elements, and two recognition sequences (Jaschke et al., 2019; Logel & Jaschke, 2020; Sanger et al., 1977; Shlomai & Kornberg, 1980). While ΦX174 has a much more complex genetic architecture than any previously AI-generated biological system, its relatively small genomic length and rich history of experimental work (Kirchberger & Ochman, 2023) also make ΦX174 a tractable and safe model for establishing whole-genome design. Notably, ΦX174 was the first complete DNA genome sequenced and synthesized (Sanger et al., 1977; Smith et al., 2003) and has continually served as a pivotal model within molecular biology (Barrell et al., 1976; Goulian et al., 1967; Jaschke et al., 2012).

Enabling generative design of complete bacteriophages first required substantial computational development of both generative models and bioinformatic prediction tools. Our full generative pipeline consisted of unsupervised pretraining with Evo 1 and Evo 2, task-specific fine-tuning on *Microviridae* genomes, prompt engineering with ΦX174-specific sequences, and inference-time guidance via predictive models of genomic architecture and host tropism. Whole-genome bacteriophage design also required substantial development of experimental methods, including a novel protocol that enabled us to screen ∼300 genome designs to yield 16 functional phages, with tropism successfully limited to the target host.

The generated phages have substantial evolutionary novelty in their sequences and structures. Some of these phages also have faster lysis kinetics than ΦX174 or directly outcompete ΦX174 in a growth assay. In a setting analogous to pathogen resistance to phage therapies, a cocktail of our generated phages rapidly overcame resistance in three different ΦX174-resistant *E. coli* strains, whereas ΦX174 alone could not overcome resistance. These results establish generative genomics as a powerful strategy for accessing novel evolutionary spaces and for potentially creating effective and evolutionarily resilient phage therapeutics.

More broadly, this study establishes a generalizable approach to generative genome design under steerable, user-specified constraints. We envision that techniques developed in this work offer a path toward designing more complex biological systems with desirable functions, potentially including the larger genomes of living organisms.

## 2. Results

### 2.1. Evo composes realistic bacteriophage genomic sequences

Genome-scale design is uniquely complex because it must reconcile features such as coding and noncoding elements, architectural arrangement, gene directionality, structural motifs, and even interactions with elements not encoded in the designed sequence (**Figure 1A**). We hypothesized that, with advances in genome language models and with the appropriate training data, it would be possible to learn underlying evolutionary constraints that could allow for the generation of phage genome sequences not yet seen in nature (**Figure 1B**).

We investigated whether Evo 1 and Evo 2, which were pretrained on large corpora of DNA sequences including over two million bacteriophage genomes (Brixi et al., 2025; Nguyen, Poli, Durrant, et al., 2024), had baseline capability for generating novel phage-like sequences (**Figure 1C**). We leveraged special taxonomic sequence labels that were included alongside genomic sequences during pretraining (Brixi et al., 2025; Nguyen, Poli, Durrant, et al., 2024) to specifically prompt the models to generate phage-like sequences, using three prompts corresponding to major viral realms: *Duplodnaviria* (double-stranded DNA viruses), *Monodnaviria* (single-stranded DNA viruses), and *Riboviria* (RNA viruses) (**Figure 1C**; **Methods**).

We first evaluated whether the generated sequences resembled natural phage sequences using geNomad, a widely used viral classification tool (Camargo et al., 2023). As a baseline, geNomad reliably classified 89–100% of natural phage sequences as viral depending on the realm (geNomad mean score > 0.96), whereas controls consisting of scrambled natural sequences were rarely classified as viral (1.9–11%, mean score > 0.73) (**Figure S1A**). In contrast, 19–33% of Evo 1 generations were classified as viral (mean score > 0.80) and 34–38% of Evo 2 generations were classified as viral (mean score > 0.87), depending on the prompt (**Figure 1D**). These results demonstrate that both pretrained models can generate phage-like sequences, with Evo 2 showing stronger baseline performance.

Querying generated sequences against the nucleotide BLAST database (Camacho et al., 2009) revealed that generated sequences are distinct from those in nature (**Figure 1E,F**). Despite their novelty, the sequences retained a high predicted coding density and genetic features reminiscent of natural phage genomes (**Figure 1F,G**). Structure prediction (Lin et al., 2023) of the proteins generated by both Evo 1 and Evo 2 yielded predicted confidence scores consistent with those expected for structure prediction of natural proteins (**Figure 1H**; **Figure S1B**). When compared to proteins in the Evo models’ pretraining data, OpenGenome (Brixi et al., 2025; Nguyen, Poli, Durrant, et al., 2024), and to proteins in the PHROGs database, a comprehensive database of phage proteins (Terzian et al., 2021), generated proteins generally showed low sequence identity with a realistic diversity of gene annotations (**Figure 1I,J**). Together, these data demonstrate that pretrained Evo models can design biologically realistic phage genomic sequences while introducing sequence diversity beyond natural evolution.

### 2.2. Generative design of novel bacteriophages with target host tropism

We next hypothesized that generative genomic design, with appropriate templates and constraints, could propose novel, complete, and viable phage genomes (**Figure 2**). To achieve bacteriophage design, we devised a workflow (**Figure 2A**) that consists of: (1) selecting a target host of biological or therapeutic interest, (2) choosing a design template—a phage genome known to infect that host—and collecting evolutionarily related sequence data, (3) training or fine-tuning a genome language model on that sequence data, (4) establishing design constraints based on the design template, (5) computationally evaluating and filtering generated sequences using those constraints, and (6) experimentally validating the designed genomes.

**Figure 2.**
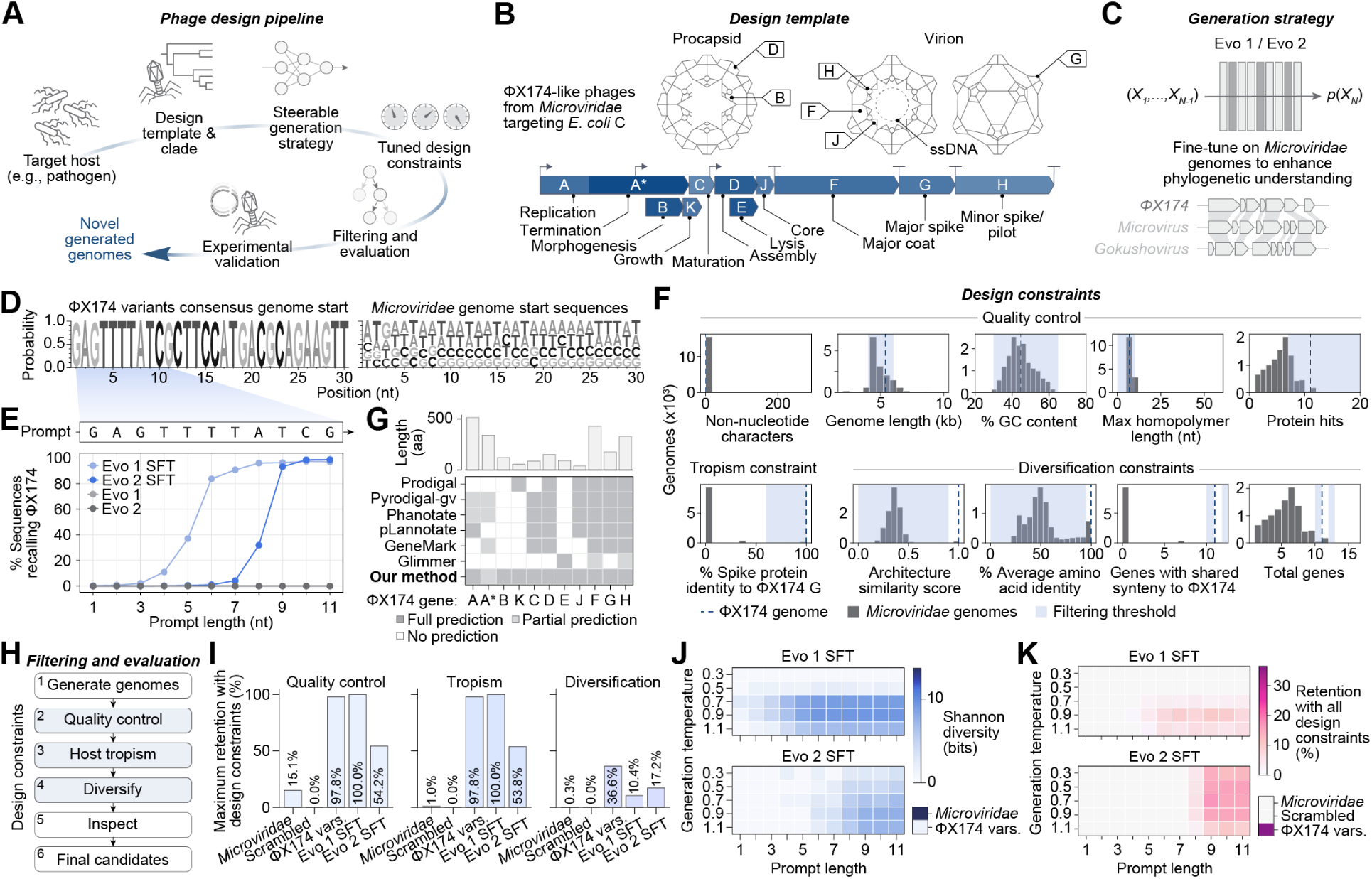
Generative design of novel bacteriophages with target host tropism. **(A)** Phage genome design workflow. **(B)** Genetic architecture of our design template, ΦX174, a *Microviridae* phage that uses *E. coli* C as a host. **(C)** For our generation strategy, we specialized Evo 1 and Evo 2 on *Microviridae* genomes through supervised fine-tuning (SFT) to enhance its ability to generate ΦX174-like sequences. **(D)** Sequence logos showing conserved nucleotides at the start of ΦX174 variant genomes compared to *Microviridae* genomes in our training data. **(E)** Increasing the number of ΦX174 nucleotides in the prompt quickly improves recall with the SFT models whereas the base models fail to recall ΦX174 across all prompt lengths. **(F)** Design constraints selected for genome filtering, with thresholds (blue) chosen against natural *Microviridae* distributions (gray) and ΦX174 (dotted line). **(G)** Benchmarking of six gene prediction methods on the genome of ΦX174 shows that overlapping genes are systematically missed, requiring us to create a new method that predicts all genes in ΦX174. **(H)** Final filtering and evaluation steps for generated genomes. **(I)** Maximum sequence retention rate across Evo 1 and 2 SFT-generated sequences after applying design constraints, with natural *Microviridae*, scrambled *Microviridae*, and ΦX174 variant (vars.) controls. The selected design constraints retain ΦX174-like sequences across quality control, tropism, and diversification filters. **(J–K)** Generated sequences have high Shannon diversity after tropism filtering **(J)** and maintain a high retention rate even after further diversification filtering **(K)**. Diversity and retention rate both increase with generation temperature and prompt length. *Microviridae*, scrambled *Microviridae*, and ΦX174 variants are shown for comparison. 𝑛 = 1000 sequences per parameter combination.

Following these steps, we selected *E. coli* C and its native phage ΦX174 as our design template (**Figure 2B**). ΦX174 is a well-studied phage that belongs to *Microviridae*, a family of phages with small genomic sizes (typically ∼4-6 kb) and an abundance of available sequencing data (**Figure S2A**) (Brister et al., 2015; Kirchberger & Ochman, 2023), while *E. coli* C is a non-pathogenic, well-characterized host strain (Michel et al., 2010). These features make ΦX174 and *E. coli* C an ideal testbed for generative phage design.

Given the success of previous design tasks that required supervised fine-tuning (SFT) to generate coherent CRISPR-Cas systems and transposable elements (Nguyen, Poli, Durrant, et al., 2024), we fine-tuned Evo 1 7B 131K and Evo 2 7B 8K on a dataset of approximately 15 thousand *Microviridae* sequences (**Figure 2C**; **Figure S2B**; **Figure S3A–F**), resulting in higher fidelity language modeling of ΦX174-like genomes (**Figure S3G,H**). We then sampled sequences from the *Microviridae* SFT models, leveraging the property that they were trained to initiate full-length genome generation from the very first position of their input context. While the genomic start sequences of all *Microviridae* sequences in the training data were highly diverse, all of the ΦX174-like genomes started with the same consensus nucleotides (**Figure 2D**). By prompting with a portion of this consensus sequence, we were able to generate ΦX174-like sequences with both Evo 1 and Evo 2 SFT models. In contrast, the base models failed to recall ΦX174 across all tested lengths of the consensus sequence prompt (**Figure 2E**).

To assess the quality of generated DNA sequences, we developed a set of genome-level design constraints by computing various statistics on natural *Microviridae* sequences, including ΦX174 (**Figure 2F**). We organized these constraints into three tiers: sequence quality, tropism specificity, and evolutionary diversity. As basic sequence quality control, we filtered out generated sequences containing non-nucleotide characters, enforced lengths between 4–6 kb and GC content within 30–65%, and excluded sequences with DNA homopolymers longer than 10 bases.

We also sought to add constraints on the protein-coding genes of our generated genomes. However, we found that none of six broadly used gene annotation tools were able to annotate all 11 genes on the wild-type ΦX174 sequence (**Figure 2G**; **Figure S4A**), reflecting the challenge of predicting overlapping open reading frames (ORFs) (**Figure S4A,B**) (Wright et al., 2022). To overcome this limitation, we built a bespoke CDS prediction method tailored to ΦX174-like sequences (**Figure S4C**; **Methods**) that was able to fully annotate all genes in ΦX174, with the exception of gene A*, which was partially predicted. With our new method, we applied an additional quality control constraint requiring at least seven predicted protein hits to natural ΦX174 proteins. Because host range is largely determined by the ability of viral spike proteins to bind host cell receptors (Michel et al., 2010; Sun et al., 2017), we applied a tropism constraint requiring that generated genomes encode spike proteins with moderately high sequence identity (≥ 60%) to the ΦX174 spike protein (**Figure 2F**).

We encouraged evolutionary novelty by introducing an optional set of diversification filters (**Figure 2F**). We preferred genomes with <95% average amino acid identity (AAI) to natural proteins, directly promoting diverged proteome sequences. We also developed a “genetic architecture” constraint to capture preservation of global gene arrangement relative to ΦX174 that we used to remove sequences that too closely resembled ΦX174 (**Figure 2F**; **Figure S5**; **Methods**). We favored genomes with 10 or 12 genes in total or those sharing synteny with 10 or 12 of ΦX174’s genes, allowing for variants with single gene losses or gains.

With our full set of design constraints, we narrowed the set of generated sequences through successive rounds of quality control, tropism filtering, and diversification filtering (**Figure 2H**). These filters also separated natural ΦX174-like sequences from other *Microviridae* genomes and scrambled *Microviridae* sequences (**Figure 2I**). After tropism filtering, we retained as much as 100% of the Evo 1 SFT generations and 53.8% of the Evo 2 SFT generations. After diversification-based filtering, we retained 10.4% of Evo 1 generations and 17.2% of Evo 2 generations.

Finally, building on our prior finding that sequence context strongly influences generative outputs (Merchant et al., 2024), we systematically assessed how prompt length alongside autoregressive sampling temperature influenced the diversity and quality of generated genomes. We found that prompting with the first nine or more nucleotides from the ΦX174 consensus sequence (**Figure 2D**) led to simple, memorized recall of ΦX174 with minimal diversity at low sampling temperatures (**Figure S6**). In contrast, prompting with only the first one or two nucleotides from the consensus sequence did not provide sufficiently strong conditioning to produce ΦX174-like generations. Notably, steering toward diverse, ΦX174-like sequences required carefully tuning the length of the consensus-sequence prompt to be around 4–9 nucleotides and the sampling temperature to be in the 0.7–0.9 range (**Figure 2J**; **Figure 2K**; **Figure S6**).

Both Evo 1 and Evo 2 SFT models produced highly diverse sequence populations even after tropism filtering, with Shannon diversity, a measure of species biodiversity (Peet, 1974), increasing alongside temperature and prompt length (**Figure 2J**). Importantly, this diversity did not come at the cost of design quality, with a high percentage of sequences being retained after applying our diversification filters (**Figure 2K**). In total, these analyses show that by combining steerable generation with systematic design constraints, Evo models can propose phage genomes that are realistic, template-guided, and evolutionarily novel.

### 2.3. Creating functional generated bacteriophage genomes

Beyond computational design, we then sought to experimentally test whether the generated genomes encode viable phages (**Figure 3**). Using our evaluation criteria, we curated a set of 302 diverse generated phage genome candidates (**Figure S7**; **Figure S8**). We designated these phages ‘Evo-Φ’ followed by a unique numeric sequence identifier. These genomes spanned 4–6 kb in length, have average amino acid identities (AAIs) to natural proteins as low as 63%, and retained spike protein sequence identities largely above 85% (**Figure 3A**). Most candidates shared more than 40% nucleotide identity with ΦX174 and genomes in the *Microviridae* training data. CheckV, a tool for assessing the quality of viral sequences, classified over 87% of the generated sequences as High Quality or Complete (Nayfach et al., 2021). Many of the generated genes also shared no coding sequence similarity with any gene in ΦX174, resulting in breaks in synteny (**Figure 3B**). Most genomes encoded 11 genes in total, with 10 preserving synteny with ΦX174 (**Figure 3C**).

**Figure 3.**
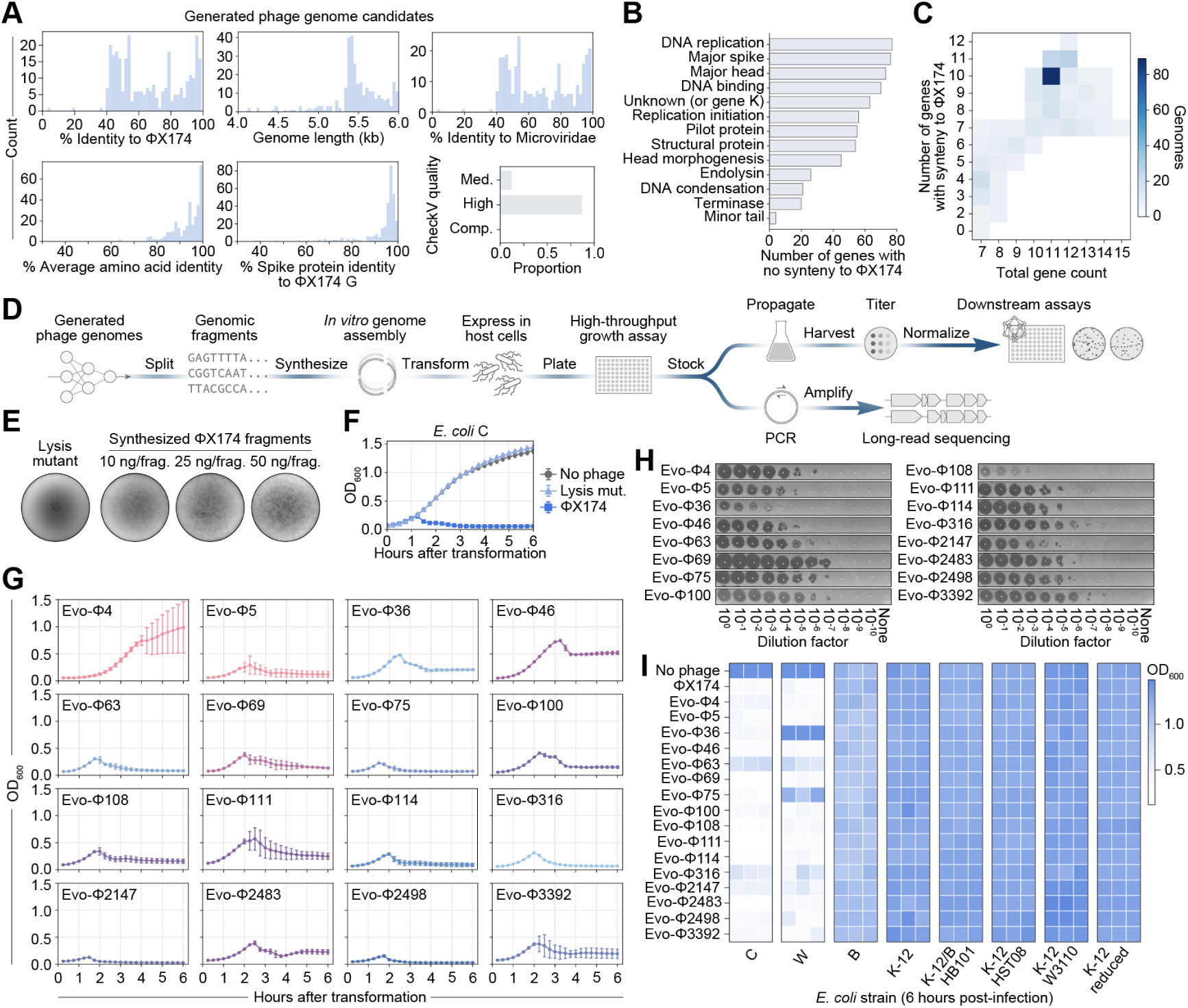
Creating functional generated bacteriophage genomes. **(A)** Final generated phage candidates meet our quality criteria while capturing abundant sequence diversity. kb, kilobase; Med., Medium; Comp. Complete. **(B)** Many generated sequences encode genes with low sequence identity to ΦX174, resulting in breaks in gene synteny. **(C)** A heatmap of total gene count versus number of syntenic genes shows that most sequences contain a single-gene break in synteny to ΦX174, balancing conservation with novelty. **(D)** Workflow for experimental validation of generated phage genomes. **(E)** *E. coli* C transformed with synthesized ΦX174 genome assemblies show that phage plaques robustly form across assembly conditions. In contrast, the same genomes but with loss-of-function mutations in lysis genes do not form plaques. ng, nanogram; frag., fragment. **(F)** Growth curves of *E. coli* C transformed with no phage (gray), ΦX174 lysis mutant (mut., light blue), or wild-type ΦX174 (dark blue) genome assembly reveal strong growth inhibition by ΦX174. Data point, mean OD_600_ value; error bar, standard deviation; 𝑛 = 3 growth replicates. **(G)** Growth curves of generated genome assemblies transformed in *E. coli* C exhibiting strong growth inhibition. **(H)** Representative titrations of propagated phage candidates. **(I)** Growth inhibition measured by OD_600_ at 6 hours after infection of *E. coli* cultures with no phage, ΦX174, or generated phages shows that ΦX174 and generated phages inhibit growth in the target strain *E. coli* C, and in *E. coli* W, but not in six other strains tested, demonstrating the robustness of tropism filtering. Each column is an infection replicate.

We developed a growth assay for testing phage replication and lysis similar to previously established phage rebooting protocols (**Figure 3D**) (Faber et al., 2019; Leuven et al., 2021). A mature ΦX174 virion contains a circular single-stranded genome that becomes double-stranded during replication (Fiers & Sinsheimer, 1962; Wickner & Hurwitz, 1974), a stage that enables phage rebooting through transformation of a circular double-stranded DNA (dsDNA) product (Faber et al., 2019; Leuven et al., 2021) that we can obtain via *in vitro* assembly of dsDNA fragments (**Figure 3D**). With this protocol, we observed that genomes of ΦX174 robustly formed plaques on a plate of *E. coli* C, whereas variants with mutated lysis genes did not (**Figure 3E**; **Figure S9A**). In our growth inhibition assay, ΦX174 impeded growth of *E. coli* C in less than two hours (**Figure 3F**). Upon successful growth inhibition, we created a stock of the phage-infected culture, verified its identity by long-read sequencing, then propagated and titrated the phage for downstream assays (**Figure S9B,C**; **Methods**).

Upon validating our screening protocol with wild-type ΦX174, we successfully synthesized and assembled 285 out of 302 generated genomes; the remaining failed due to high-complexity DNA synthesis. Measuring bacterial growth inhibition as an indication of phage viability, we observed 16 generated phage transformations that inhibited growth of *E. coli* C (**Figure 3G**; **Figure S10A**). When transformed in *E. coli* K-12, none of the 285 assemblies resulted in growth inhibition, supporting the robustness of the tropism constraint filter (**Figure S10B**). Sequence-verification of the 16 candidates revealed nine genomes with no acquired mutations, whereas the other seven acquired some single nucleotide variants (SNVs) or deletions with respect to the synthesized sequences (**Figure S11**). The generated phages also showed variable titers when propagated (**Figure 3H**; **Figure S12**).

Finally, we tested the host range of the generated phages across eight *E. coli* strains, including *E. coli* C, *E. coli* B, *E. coli* W, and five variants of *E. coli* K-12 (**Figure 3I**; **Figure S13**). Surprisingly, ΦX174 and 15 out of 16 of the generated phages could also inhibit growth in *E. coli* W, a host not previously associated with ΦX174 (Archer et al., 2011; Michel et al., 2010), although with much more variation in growth kinetics (**Figure S13**). We did not observe growth inhibition in the other six *E. coli* strains, suggesting that the generated phages maintain a high specificity for the intended host, *E. coli* C. More broadly, these results demonstrate that genome language models, combined with inference-time steering and filtering, can design viable phage genomes.

### 2.4. Generated bacteriophages reveal sequence and structural insights

Having validated the viability of our generated phages, we next examined the extent of their evolutionary novelty (**Figure 4**). Upon analyzing the mutational differences within the generated genomes relative to ΦX174 and other *Microviridae* phages (**Figure 4A–D**), we observed hundreds of synonymous, nonsynonymous, and noncoding mutations (**Figure 4A**) that include:

- a novel gene J insertion in Evo-Φ63;
- extended noncoding regions in Evo-Φ63, Evo-Φ2147, Evo-Φ2483, and Evo-Φ2498;
- loss of gene K in Evo-Φ114;
- large, putative truncations of gene C in Evo-Φ75 and Evo-Φ100, and of gene B in Evo-Φ316;
- large, putative elongations of gene E in Evo-Φ4 and Evo-Φ46; and
- swapping of gene J in Evo-Φ36 with gene J from Escherichia phage G4, a swap previously found non-viable for wild-type ΦX174 (Fane et al., 1992; Ogunbunmi et al., 2021; Roznowski et al., 2020).

**Figure 4.**
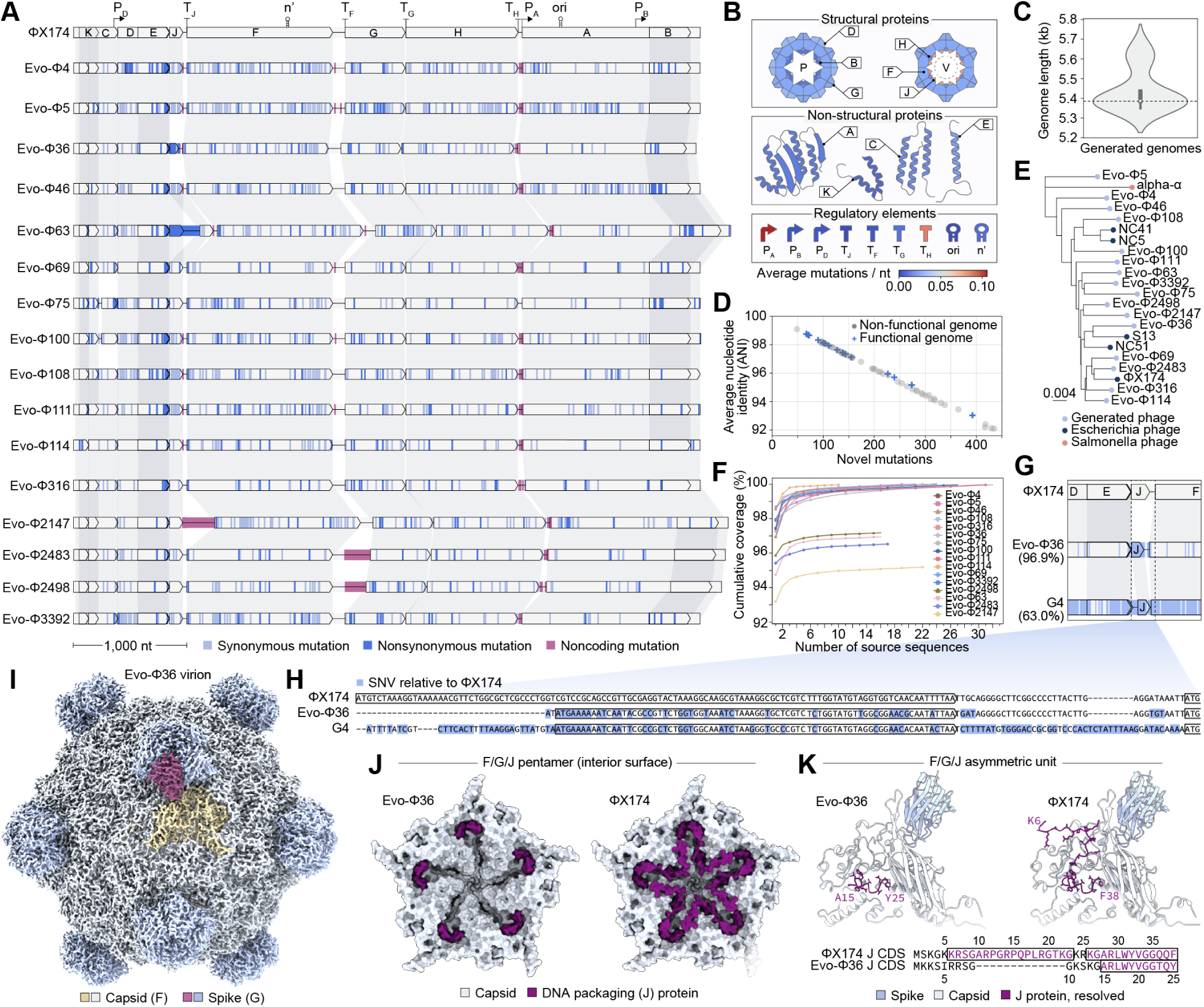
Generated bacteriophages reveal sequence and structural insights. **(A)** Synteny plot of ΦX174 and functional generated bacteriophages, highlighting hundreds of synonymous (light blue), nonsynonymous (dark blue), and noncoding (red) mutations compared to ΦX174. Genomes are in order by name. **(B)** Average nucleotide (nt) mutational frequencies, normalized to length, of structural proteins, non-structural proteins, and regulatory elements across the generated phage genomes show that gene J and regulatory elements are mutational hotspots. **(C)** Generated genomes exhibit a range of lengths. Dotted line, genome length of ΦX174; white dot, median; gray box, interquartile range (IQR); whiskers, 1.5× IQR. **(D)** Percent sequence identity and number of novel mutations of functional (blue) and non-functional (gray) generated sequences compared to their top nucleotide BLAST hit in the *Microviridae* training data. **(E)** Neighbor-joining phylogenetic tree of functional generated phages (light blue) and representative *Microviridae* phages (dark blue and pink). **(F)** Percent cumulative sequence coverage of generated phages highlights that mutations in most generated phages cannot be completely attributed to mutations seen in nature. The sequences were aligned to sequences by nucleotide BLAST in the core_nt database until all nucleotides were accounted for or there were no significant hits for remaining nucleotides. **(G–H)** Synteny plot **(G)** and detailed view of gene J **(H)** and its surrounding intergenic regions of ΦX174, Evo-Φ36, and phage G4, with single nucleotide variations (SNVs) compared to ΦX174 highlighted in blue. **(I)** Cryo-EM density map of Evo-Φ36 virion, highlighting individual subunits of the spike (pink) and capsid (yellow). Remaining spikes (light blue) and capsids (gray) are shown. **(J)** Interior surface view of the capsid (F, gray) and spike (G, not visible) pentamers of Evo-Φ36 (left) and ΦX174 (right), with their cognate J proteins (purple) reveals distinct capsid interactions and putative genome packaging modes between the two phages as modeled into the cryo-EM density map. **(K)** Asymmetric units including F, G, and J of Evo-Φ36 (left) and ΦX174 (right) show resolved residues of J.

The most mutated elements amongst the 16 genomes were promoter A, terminator H, and gene J, with average mutation rates of 0.11, 0.09, and 0.07 mutations per nucleotide, respectively (**Figure 4B**). The regulatory elements of Terminator J and the origin of replication (ori) were the least mutated, with no mutations across all sixteen genomes. The generated genomes also had a range of lengths, from 99% to 105% of the length of ΦX174 (**Figure 4C**).

When we compared the generated genomes to *Microviridae* sequences in the training data, they contained between 67 and 392 novel mutations, with nucleotide sequence identities between 93.0% and 98.8% (**Figure 4D**). The nearest natural genomes to the generated genomes consisted of several *Microviridae* phages that infect *E. coli*, including ΦX174, NC41, NC5, NC51, and S13 (**Figure 4E**). Evo-Φ2147 has 392 mutations—representing 93.0% average nucleotide identity (ANI)—with respect to its nearest natural genome (**Figure 4D**), phage NC51; notably, natural genomes with less than 95% ANI to any known phage would typically qualify as a new species (Turner et al., 2021).

Interestingly, over 50 generated sequences that were not viable in our growth inhibition screens contained a comparable number or fewer mutations than the viable generated phages when aligned to their nearest natural genomes, highlighting the difficulty of designing functional mutations at the genome-scale. Further, mutations in 13 of the generated genomes could not be recapitulated from any known natural sequences (**Figure 4F**; **Figure S14**), including large portions of the novel noncoding regions in Evo-Φ63, Evo-Φ2147, and Evo-Φ2483. Together, these results demonstrate the ability of genome language models to design complete genomes with high sequence novelty.

Beyond genetic diversity, we examined structural novelty in the generated phages. In Evo-Φ36, gene J, a genome-packaging protein that also supports the capsid (Bernal et al., 2004), was replaced with a homologous protein found in phage G4, differing by four synonymous SNVs. Yet, the J–F intergenic region remained more similar to ΦX174 (**Figure 4G,H**). G4 is a distantly related Microvirus with only 63.0% genome identity to ΦX174 (**Figure S15A**) (Godson et al., 1978), whereas Evo-Φ36 shares 96.9% identity with ΦX174. The G4 J protein is 25 amino acids long compared to 38 amino acids in ΦX174, lacking portions of domains 0 and I that contribute to capsid binding in ΦX174 (**Figure S15B**) (Bernal et al., 2004; Ogunbunmi et al., 2021). Interestingly, prior work has shown that swapping G4 J into ΦX174 is not viable (Fane et al., 1992; Ogunbunmi et al., 2021; Roznowski et al., 2020). Remarkably, Evo-Φ36 is viable despite encoding the G4 J, highlighting Evo’s ability to design genomes that integrate context-dependent structural compatibility.

Given the unusually small J protein in Evo-Φ36, we sought to understand its structural consequences. AlphaFold 3 revealed strikingly different orientations of J in the capsid pentamers of Evo-Φ36 compared to ΦX174 and G4 (**Figure S15C–E**) (Abramson et al., 2024). To further explore these interactions, we solved the structure of ΦX174 to a resolution of 2.8 Å and Evo-Φ36 to a resolution of 2.9 Å by cryogenic electron microscopy (cryo-EM) (**Figure 4I**; **Figure S16**; **Figure S17**; **Figure S18**; **Table S1**). Consistent with previous findings (McKenna et al., 1992), we observed that the hydrophobic C-terminus of the J protein in ΦX174, domain II, interacts with the interior surface of the capsid, while the more basic domains 0 and I tether the genomic DNA while structurally supporting the capsid (**Figure 4J,K**; **Figure S19**). Despite introducing more polarity at the capsid-J interface by substitution of the C-terminal F→Y, Evo-Φ36 J maintains a similar overall binding mode at its C-terminus (**Figure 4K**). Notably, we could not resolve 14 residues at the N-terminus of Evo-Φ36 J, whereas the corresponding residues in ΦX174 J visibly interact with the capsid. This difference suggests that the Evo-Φ36 N-terminus is likely unstructured with respect to the capsid and primarily functions in DNA binding. Taken together, these findings show that despite divergent sequence contexts, Evo-Φ36 J preserves a compatible interaction with its capsid, denoting how generative design can uncover novel protein-protein co-evolutionary solutions.

### 2.5. Generated bacteriophages exhibit high fitness

Generative genomics can propose novel genetic sequences, including large-scale sequence changes, that are not constrained by natural selection pressures. We therefore hypothesized that functional mutations in the generated phage genomes might confer fitness advantages relative to wild-type ΦX174 (**Figure 5**). We competed ΦX174 and all generated phages against each other in a single *E. coli* C population by co-infecting the 16 generated phages and ΦX174 at equal multiplicity of infection (MOI) and measuring the cumulative fold change (FC) in sequencing read counts of each phage over time (**Figure 5A**). At the end of a competition, phages with the highest cumulative FC are those that had the largest relative increase in their own population size, indicating high fitness.

**Figure 5.**
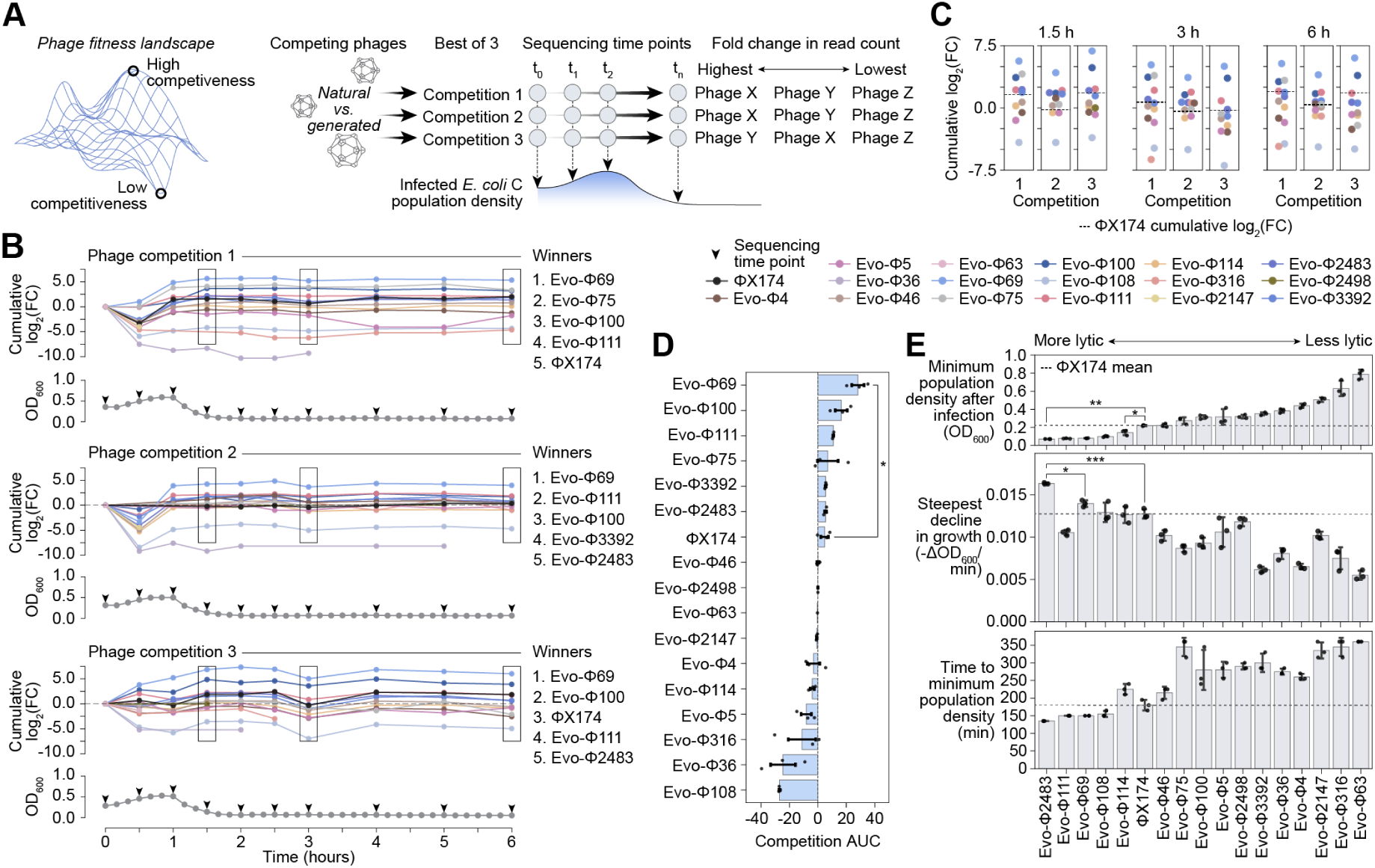
Generated bacteriophages exhibit high fitness. **(A)** Phage fitness competition assay workflow. **(B–C)** In three competitions, generated phages and ΦX174 competed head-to-head in *E. coli* C at equal multiplicity of infection (MOI). We tracked cumulative fold change (log_2_(FC)) of sequencing read counts over six hours **(B)**. Many generated phages matched or surpassed ΦX174’s performance at various time points **(C)**, indicating a higher relative fitness. Growth curves of infected *E. coli* C populations show corresponding suppression of bacterial growth. Rectangular boxes, enlarged plots in **(C)**; arrowheads, sequencing sample extraction time points. Dotted line, cumulative log_2_(FC) of ΦX174. **(D)** Area under the curve (AUC) of the cumulative log_2_(FC) of phage read counts shows that generated phages outcompeted ΦX174 over the whole time course. Statistical significance was determined by one-way ANOVA with Tukey HSD (*𝑝-adj < 0.05). Bar height, mean; error bar, standard deviation; circles, 𝑛 = 3 competitions; dotted line, AUC of 0. **(E)** Growth dynamics of *E. coli* C infected with generated phages and ΦX174 individually show that several generated phages exhibit lower minimum population density after infection, steeper decline in host growth rate, and shorter time to minimum population density, together indicating stronger lytic capabilities. Statistical significance was determined by one-way ANOVA with Tukey HSD (*𝑝-adj < 0.05; **𝑝-adj < 0.01; ***𝑝-adj < 0.001). Bar height, mean; error bar, standard deviation; circles, 𝑛 = 3 infections; dotted line, mean value of ΦX174.

We observed that three generated phages, Evo-Φ69, Evo-Φ100, and Evo-Φ111, appeared in the top five phages at the end of three independent competition experiments (**Figure 5B–D**). ΦX174 only appeared in the top five phages in competitions 1 and 3, at most ranking in third place. Remarkably, in all three competitions, Evo-Φ69 outcompeted all other phages, with cumulative fold changes between 16× and 65× after six hours of infection. In contrast, at the same time point, ΦX174’s cumulative fold change ranged from 1.3× to 4.0× from its initial infection count.

We next examined the infection dynamics of each phage by measuring how quickly and strongly each drove the host population to its minimum density, focusing on the timing, rate, and depth of lysis (**Figure 5E**; **Figure S20**). We found that one phage, Evo-Φ2483, exhibited the fastest and strongest lytic capabilities. On average, Evo-Φ2483 drove the host population to a minimum OD_600_ of 0.07, with a maximum growth rate decline of –0.02 OD_600_/min, reaching this minimum in 135 minutes; by comparison, ΦX174 infections resulted in a higher minimum OD_600_ of 0.22, a slower maximum rate of decline of –0.01 OD_600_/min, and 180 minutes to reach minimum density. Several other candidates—Evo-Φ111, Evo-Φ69, Evo-Φ108, and Evo-Φ108—also reached significantly lower host population densities than ΦX174. Notably, Evo-Φ2483 only ranked fifth in two of the three competition assays (**Figure 5B**), underscoring that lytic effect alone does not determine overall fitness. As a whole, these results support the ability of genome language models to design high fitness mutations at the genome-scale, yielding diverse phenotypic outcomes for phage life cycles which could benefit phage-based biotechnologies.

### 2.6. Generated bacteriophages rapidly overcome bacterial resistance

Phage therapy is emerging as a promising alternative to antibiotics, but its effectiveness can be limited by rapid evolution of resistant bacteria (Strathdee et al., 2023). We hypothesized that the diverse phages produced by our genome design method could form a cocktail that more readily overcomes bacterial resistance (**Figure 6**) (Kim et al., 2025; Pires et al., 2016; Pirnay, 2020). To investigate our hypothesis, we first evolved three ΦX174-resistant *E. coli* C cultures (**Figure 6A**; **Figure S21A**; **Methods**). Whole-genome sequencing of three isolated strains from each resistant *E. coli* C culture revealed that each strain independently developed novel mutations within the *waa* operon (**Figure 6A**; **Figure S21B**), which is associated with LPS synthesis. We called these strains CR1, CR2, and CR3. In particular, strain CR1 contained a missense mutation, L259W, in the *waaT* gene; strain CR2 contained a single base deletion at nucleotide position 485 of *waaT*, resulting in a premature stop codon followed by potential reinitiation at a downstream in-frame start codon; and strain CR3 contained a missense mutation, A128D, in the *waaW* gene. Mutations in *waaT* and *waaW* have previously been observed to confer resistance against ΦX174 (Romeyer Dherbey et al., 2023), suggesting that strains CR1, CR2, and CR3 likely exhibit resistance through modifications to LPS synthesis.

**Figure 6.**
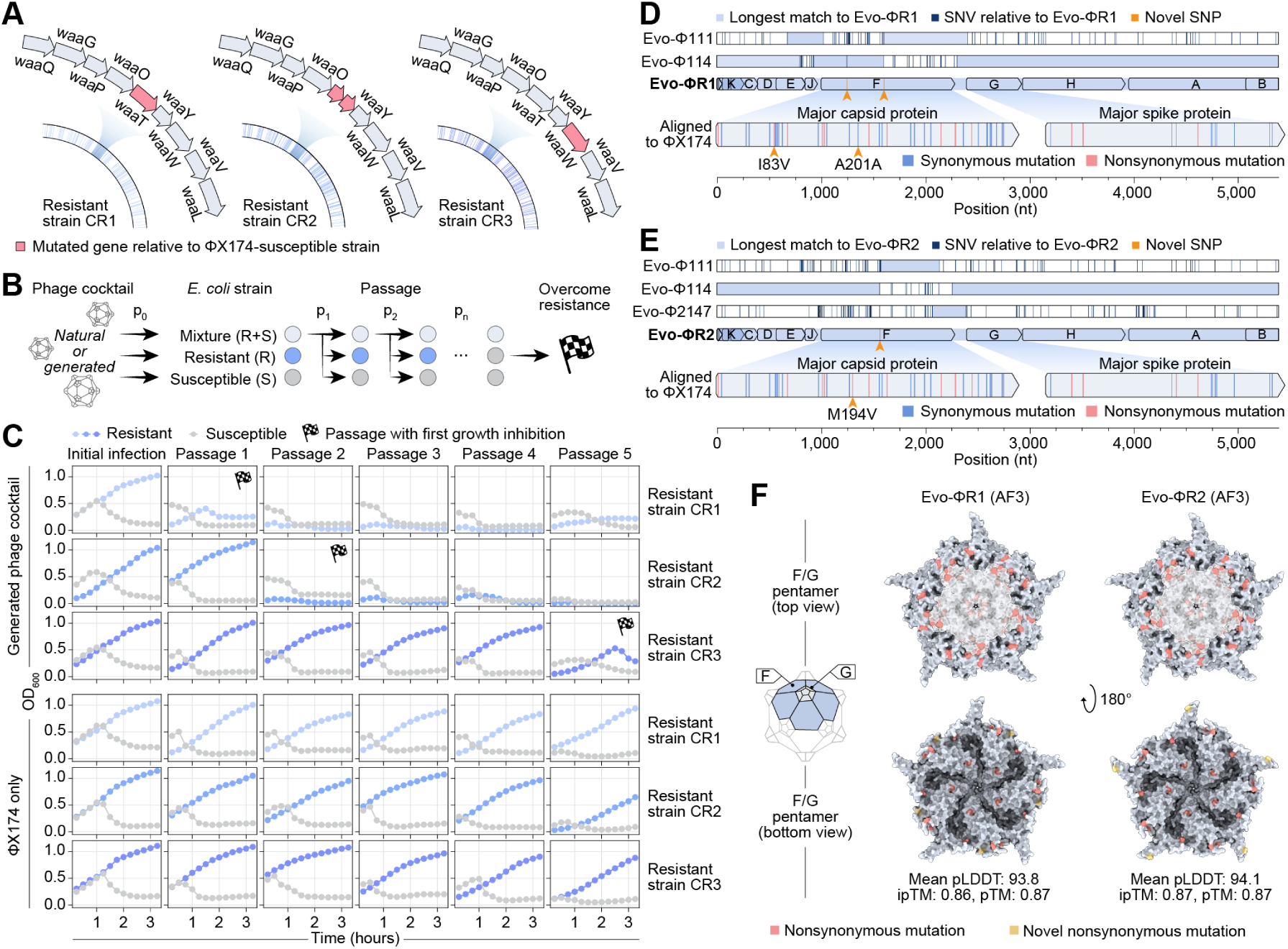
Generated bacteriophages rapidly overcome bacterial resistance. **(A)** Whole-genome sequencing of three ΦX174-resistant *E. coli* C strains revealed mutations in the *waa* operon absent in susceptible *E. coli* C, which functions in lipopolysaccharide synthesis. **(B)** Experimental setup for evolving phage counter-resistance by serially passaging cocktails of generated phages and ΦX174, or ΦX174 alone, on susceptible and resistant *E. coli* C. **(C)** Growth curves show that ΦX174 alone fails to overcome resistance, whereas generated phage cocktails suppress growth of all resistant cultures within five passages. Checkered flag, first passage with growth inhibition. **(D–E)** Alignments of generated phages against the predominant resistant phages Evo-ΦR1 **(D)**, Evo-ΦR2 **(E)** capable of infecting resistant strain 1 and 2, respectively, show that they are derived from generated genomes. Generated phages used in the alignments are those with the longest identical sequences (light blue) without single nucleotide variations (SNVs; dark blue) to each resistant phage such that they collectively minimize the number of novel mutations (yellow) observed in the resistant phage. Major capsid and spike proteins of each resistant phage aligned to ΦX174 major capsid and spike proteins are below, with synonymous (blue) and nonsynonymous mutations (pink) relative to ΦX174. **(F)** AlphaFold 3 (AF3) predictions of capsid (light blue) and spike (light gray) pentamers show that most capsid and spike mutations appear on the exterior of resistant phages. Nonsynonymous and novel nonsynonymous mutations relative to ΦX174 are highlighted in pink and yellow, respectively. pLDDT, predicted local distance difference test score; ipTM, interface predicted template modeling score; pTM, predicted template modeling score.

Next, we explored whether ΦX174 only, or a mixture of all 16 generated phages and ΦX174, could overcome resistance in strains CR1, CR2, and CR3. We challenged three separate cultures: a ΦX174-susceptible strain, a ΦX174-resistant strain, and a mixture of the two strains to create a selective pressure for phages replicating in susceptible cells to evolve the ability to infect resistant cells (**Figure 6B**). After each challenge, we passaged the supernatant from the mixed strains into fresh cultures and observed their growth dynamics.

Upon initial infection, the generated phage cocktail and ΦX174 successfully inhibited growth of the susceptible strains but did not inhibit growth of the resistant strains (**Figure 6C**). Strikingly, the generated phage cocktail was able to successfully inhibit growth of strain CR1 after a single passage, CR2 after two passages, and CR3 after five passages. In contrast, ΦX174 alone could not inhibit growth of any resistant strains even after all five passages.

We isolated and sequenced several individual phages that successfully grew on strains CR1 and CR2, which revealed a single predominant phage genome responsible for overcoming each resistant strain (**Methods**), which we designated Evo-ΦR1 and Evo-ΦR2, respectively. Sequence alignment revealed that all of Evo-ΦR1’s genome could be collectively composed from five segments of Evo-Φ111 and Evo-Φ114, with the exception of two novel SNVs in the major capsid protein (**Figure 6D**). All of Evo-ΦR2’s genome could be composed from four segments of Evo-Φ111, Evo-Φ114, and Evo-Φ2147, with the exception of a single novel SNV in the major capsid protein (**Figure 6E**). The ability to overcome resistance likely arose from recombination and acquired mutation events involving two or three of the Evo-generated phages.

Given the resistant mutations in the LPS gene synthesis operons of strains CR1 and CR2, we speculated that key mutations in the predominant resistant phages likely decorated the outer surfaces of their major capsid and spike proteins, since these proteins modulate LPS binding (Michel et al., 2010). Indeed, for both Evo-ΦR1 and Evo-ΦR2, we observed 15 missense mutations across the capsid and spike proteins that were not present in ΦX174, 14 of which were generated by Evo (Fig. 6D,E). The majority of the mutations appear on the outer surface of the virions, as predicted by AlphaFold 3 (**Figure 6F**; **Figure S21C,D**) (Abramson et al., 2024). These data show that evolutionary innovations generated by Evo likely contributed to resilience against bacterial resistance and, more broadly, suggest the utility of generative models for producing genetically diverse phage cocktails that could translate into improved therapeutic efficacy.

## 3. Discussion

In this work, we leveraged genome language models to achieve the first generative design of complete bacteriophage genomes. We established a computational framework for specifying our design goals, including the development of a new gene annotation method and diverse scoring metrics, allowing us to controllably design toward a target genomic architecture and host tropism. In particular, our design template was based on ΦX174, a tractable, safe, and historically significant model genome (Barrell et al., 1976; Goulian et al., 1967; Kirchberger & Ochman, 2023; Sanger et al., 1977; Smith et al., 2003). We systematically evaluated thousands of computationally generated sequences and experimentally tested nearly 300 designs, resulting in 16 viable phages containing substantial evolutionary diversity and enabling a phage cocktail that rapidly overcame bacterial resistance. Multiple generated phages exhibited increased fitness or faster lytic dynamics relative to ΦX174, demonstrating the ability of generative models to efficiently evolve high-fitness genomes.

Synthetic genomics has historically relied on directed evolution, random mutagenesis, or rational engineering (Coradini et al., 2020; James et al., 2024). These approaches have been limited in the scope of their achieved evolutionary novelty due to the complexity of genome sequences and the limited throughput of methods for genome editing and synthesis. Rational engineering is further limited by incomplete human understanding of biology; for example, previous systematic efforts have struggled to increase features such as phage lysis rate or genome length (Aoyama & Hayashi, 1985; Endy et al., 2000; Russell & Müller, 1984). In contrast, our approach enabled designs with substantial novelty in both nucleotide and protein sequence, including the genome of Evo-Φ63, that is 5% (268 bp) longer than ΦX174; Evo-Φ69, which outcompeted ΦX174 in competition assays; and Evo-Φ2483, which exhibited significantly faster lysis rates. Notably, the genome Evo-Φ2147 achieves a level of nucleotide sequence novelty (<95% ANI) on par with that achieved by natural evolution when producing new bacteriophage species (Turner et al., 2021).

The capability to generate novel genomes with AI systems also raises important biosafety considerations. In line with longstanding precedent for conducting biological research within established biosafety levels (Berg et al., 1975), we performed all experiments at the biosafety level appropriate for research with bacteriophages and their non-pathogenic bacterial hosts, alongside supplementary precautions (**Methods**). These established biosafety systems can be effectively adapted and applied to the generative design of new biological systems, especially when, as in this work, designs are constrained by well-characterized natural genomes as templates. Moreover, as we have previously demonstrated, the generative models themselves can possess inherent safeguards based on their training data; for instance, we have previously shown that data exclusions successfully prevent the Evo 2 models from designing eukaryotic viruses, including pathogenic human viruses (Brixi et al., 2025). We have also provided additional details in a supplementary **Biosafety and biocontainment discussion**. By continuing to build upon robust safety frameworks, the field can responsibly unlock the potential of generative models to access and engineer complex biological functions for the benefit of science and society.

We envision that the further development of generative genomics for whole-genome design will require additional methodological innovations and improved training datasets. Future approaches could leverage techniques for improving and accelerating model conditioning beyond SFT alone (Hu et al., 2021; Lewis et al., 2020; Mnih et al., 2015; Widatalla et al., 2024). Continual genomic and metagenomic sequencing projects could also contribute additional training data. Designing larger phages will pose additional challenges, particularly in cost-effective DNA synthesis and assembly. Techniques for multi-fragment genome assembly, combinatorial oligonucleotide pools, and *in vitro* transcription–translation systems are promising avenues for realizing more complex genome design (Faber et al., 2019; Levrier et al., 2024; Pryor et al., 2022).

Beyond biotechnological or therapeutic utility, generative design of whole genomes offers unique opportunities for studying evolution (Shaer Tamar & Kishony, 2022). By systematically sampling from learned genomic distributions, it is possible to explore large mutational landscapes and investigate the genetic basis of properties such as host tropism. Indeed, we observed subsequent recombination and mutation of our generated phages that likely conferred resilience against host resistance. The ability to rapidly design phage genomes tuned for host range, fitness, and resistance evasion may expand biotechnological toolkits and transform phage therapy pipelines, enabling more adaptive and resilient antimicrobial strategies (Kim et al., 2025; Mutalik & Arkin, 2022; Pirnay, 2020; Wimmer et al., 2009).

In total, our results demonstrate that generative AI can capture an underlying evolutionary design space with enough fidelity to produce novel functional bacteriophage genomes. Continued progress will likely bring about more generalizable models capable of designing across diverse biological systems with desirable functional properties. The rapid pace of improvement in generative biology suggests a future where genome design could become a core biotechnology alongside genome sequencing, synthesis, and editing, possibly enabling the generation of complete living organisms.

## Acknowledgements

We thank Brandon Ameglio, Paul Bollyky, Ashir Borah, Collin Chiu, Seyone Chithrananda, Drew Endy, Simone Evans, Patrick Hsu, Santiago Mille-Fragoso, Brian Plosky, Jessica Sacher, and Sebastián Somolinos for helpful discussions and support with the manuscript. We thank Jerome Ku and Jeremy Sullivan for assistance with computational infrastructure. We thank Twist Bioscience for support with DNA synthesis. We thank Arc Institute LabOps for assistance with experimental setups. We thank Ashir Borah, Collin Chiu, Joseph Noh, Alex Gao’s lab, Patrick Hsu’s lab, and Lingyin Li’s lab for generous sharing of materials and reagents. We thank Joseph Noh, Haoqing Wang, and the Stanford University Cryo-Electron Microscopy Center (cEMc) for assistance with cryo-EM. S.H.K., A.T.M., and G.B. acknowledge funding support from the National Science Foundation Graduate Research Fellowship Program. D.B.L. acknowledges funding support from the Fannie and John Hertz Foundation. A.T.M. acknowledges funding support from the Knight-Hennessy Graduate Scholarship Fund. B.L.H. acknowledges funding support from Arc Institute, the Gates Foundation, Stanford Institute for Human-Centered Artificial Intelligence (HAI) Hoffman-Yee Research Grants, V. Gupta, and R. Tonsing.

## 4. Author contributions

S.H.K. and B.L.H. conceived and designed the study. B.L.H. supervised the project. S.H.K. and B.L.H. collected the training data. S.H.K., G.B., and B.L.H fine-tuned the models. S.H.K. and D.G. performed the phage genome generation and evaluation. S.H.K., C.L.D., and D.G. performed the phage genome assembly transformation assays. S.H.K. and C.L.D. performed the phage sequencing, propagation, titering, and host tropism assay. S.H.K. and A.T.M. performed the phylogenetic analysis. S.H.K., C.L.D., and D.B.L. performed the cryo-EM experiments. D.B.L. processed the cryo-EM data. S.H.K., D.B.L., and M.E.W. analyzed the cryo-EM data. S.H.K. and A.T.M. performed the phage competition and bacterial resistance assays. S.H.K. and B.L.H. wrote the initial draft of the manuscript. All authors contributed to writing the final version of the manuscript.

## 6. Data and code availability

The raw and processed *Microviridae* datasets used for supervised fine-tuning are available for download at https://doi.org/10.5281/zenodo.17101843 Code for generation and fine-tuning with Evo 1 is available at https://github.com/evo-design/evo/. Code for generation and fine-tuning with Evo 2 is available at https://github.com/arcinstitute/evo2. Code for phage genome design and analysis are available at both repositories. The fine-tuned *Microviridae* model for Evo 1 is available at https://huggingface.co/evo-des ign/evo-1-7b-131k-microviridae, and the fine-tuned Microviridae model for Evo 2 is available at https://huggingface.co/evo-design/evo-2-7b-8k-microviridae. The cryo-EM density maps and resulting models for ΦX174 and Evo-Φ36 have been deposited in the Electron Microscopy Data Bank (EMDB) with EMDB codes XXXX and XXXX, and the Protein Data Bank (PDB) with PDB codes XXXX and XXXX, respectively.

## Supplementary material

### **A.** Biosafety and biocontainment discussion

The generative design of entire genomes represents a milestone in our ability to engineer biological systems, promising new biotechnologies and therapeutics. However, this powerful capability advance necessitates a thoughtful consideration of biosafety to ensure responsible development. In this study, our approach to biosafety and biocontainment leveraged computational safeguards inherent to our models, experimental design choices, and established laboratory containment protocols.

A primary layer of safety is built directly into the genome language models themselves, as the generative capabilities of these models are fundamentally shaped by their training data. In this and previous work, we have deliberately withheld all viruses with eukaryotic hosts, including those pathogenic to humans, from the models’ training data. As we have demonstrated previously, this data exclusion strategy successfully prevents Evo 2 from prediction and design tasks related to human viruses (Brixi et al., 2025).

For this study, we further specialized the models by fine-tuning them exclusively on a curated dataset of *Microviridae* genomes, a family composed entirely of bacteriophages. This targeted training means the model retains its poor performance on eukaryotic viruses while enhancing its ability to generate sequences within our specific phage design space. We have also shown that both pretraining and fine-tuning were necessary for coherent generation of complex systems such as CRISPR-Cas operons (Nguyen, Poli, Durrant, et al., 2024). Because Evo 1 and Evo 2 were not pretrained on eukaryotic viral genomes, enabling the generation of such viruses would require a substantially greater investment in terms of data, computational resources, and methodological development. While training genome language models on sequences from human viruses could have substantial utility, such as in predicting aspects of viral evolution or designing viral vectors for gene therapy, we have not taken this step prior to careful deliberation within the scientific community.

In addition to these computational safeguards, our experimental framework was designed to be intrinsically safe and controllable. We selected the lytic phage ΦX174, as well as its non-pathogenic host, *E. coli* C, as our design template. This system is well-studied, tractable, and has a long history of safe use in molecular biology research (Barrell et al., 1976; Goulian et al., 1967; Jaschke et al., 2012; Sanger et al., 1977; Smith et al., 2003). Our method also allows for precise, user-specified constraints including a strict tropism filter to ensure that generated phages were designed to specifically target our laboratory host strain, which offers more control than traditional “phage hunting” methods that isolate phages from the environment with unknown host ranges or phenotypes. The success of our tropism constraint was validated experimentally, as none of the 285 tested assemblies showed activity against off-target *E. coli* K-12 strains.

All experiments also leveraged decades of precedent and established biosafety protocols for working with phage and non-pathogenic bacterial hosts (Berg et al., 1975). As an additional precaution, all work involving phages was performed within a biosafety cabinet with appropriate PPE and dedicated equipment, which was regularly sterilized with 70% ethanol, 10% bleach, and UV treatment, and any waste was disposed of as biohazardous waste. Furthermore, the risk of significant ecological disruption by any designed ΦX174-variant phage is minimal, given both our containment strategies and the powerful systems that bacteria have evolved for phage defense.

We implemented multiple safeguards to ensure that the establishment of whole-genome design in this study was conducted responsibly. We view bacteriophages as a biologically safe testbed for developing this foundational technology, which is fundamentally distinct from the development of viruses with eukaryotic hosts. Future work leveraging AI tools to design or adapt components of human pathogens or infectious agents should be approached with substantial caution and conducted in accordance with all applicable policies, regulations, and best practices. We conclude by noting that the ability to understand and manipulate complex biological systems to cure disease and ease human suffering has been a longstanding goal of scientific research. If done safely and responsibly, generative AI systems have immense promise for advancing humanity toward this goal.

### **B.** Methods

#### **B.1.** Generative modeling of bacteriophage genomes

##### ***B.1.1.*** Generating sequences with Evo

Sequence sampling was performed using a standard autoregressive sampling algorithm with the sampling code from https://github.com/evo-design/evo/ and https://github.com/arcinstitute/evo2, which leverages kv-caching of Transformer layers (Chang & Bergen, 2024) and the recurrent formulation of Hyena layers for efficient, low-memory autoregressive generation (Brixi et al., 2025; Nguyen, Poli, Durrant, et al., 2024; Nguyen, Poli, Faizi, et al., 2024).

##### ***B.1.2.*** Generating sequences with bacteriophage realm prompts

For Evo 1 7B 131K, approximately 10,000 sequences of length 6,000 were sampled with temperature = 0.7, top-k = 4, and top-p = 1 for each prompt. Sequences were generated using the prompts |r Duplodnavi ria;k , |r Monodnaviria;k , and |r Riboviria;k . For Evo 2 7B 1M, approximately 1,000 sequences of length 6,000 were sampled with temperature = 0.7, top-k = 4, and top-p = 1 for each prompt. Sequences were generated using the prompts |r Duplodnaviria;;;;;;;|, |r Monodnaviria;;;;;;;|, and |r Riboviria;;;;;;;|.

##### ***B.1.3.*** Generated bacteriophage realm sequence analyses

For use as positive controls, natural sequences corresponding to the realms *Duplodnaviria*, *Monodnaviria*, and *Riboviria* were extracted from the OpenGenome stage 2 training dataset (Nguyen, Poli, Durrant, et al., 2024) by filtering entries whose taxonomy strings began with |r Duplodnaviria, |r Monodnaviria, or |r Riboviria, respectively. From each realm, 10,000 sequences were randomly sampled. Sequences containing only canonical nucleotides (A, C, G, T) were retained for downstream use. For use as negative controls, scrambled natural sequences were generated by randomly permuting the nucleotide order of each sequence using a fixed random seed (42) to preserve overall nucleotide composition while ablating higher-order structure.

To analyze novelty of generated sequences, the generated sequences were searched against natural sequences in the core_nt database by nucleotide BLAST (blastn, E-value = 0.5, default settings) (Camacho et al., 2009). Query coverage and percent identity of the top 100 target hits were collected for each generated sequence. To determine the proportion of generated sequences that classify as viral, the sequences were analyzed by geNomad version 1.8.0 (Camargo et al., 2023). Coding density was determined by predicting ORFs using Prodigal version 2.6.3 (Hyatt et al., 2010) and dividing the total sum of ORF lengths per sequence with the total length of the sequence. The pTM and mean pLDDT scores of predicted proteins were extracted from their structures folded by ESMFold (Lin et al., 2023), specifically the Hugging Face implementation of the facebook/esmfold_v1 checkpoint accessed via the Transformers library. Phage-like coding sequences were predicted with our gene annotation method (see ORF calling and gene annotation), and visualized by LoVis4u with flags -hl, --set-category-colour, -cA4p2, -alip (Egorov & Atkinson, 2025). Percent protein sequence identities of the generated proteins were determined by aligning them against Prodigal-predicted proteins in OpenGenome (Nguyen, Poli, Durrant, et al., 2024), and proteins in the PHROGs database (Terzian et al., 2021) with MMseqs2 version 13.45111 (Steinegger & Söding, 2017) with sensitivity = 4.0. Functional annotations curated for PHROGs (Terzian et al., 2021) were extracted for generated proteins with significant alignments against proteins in the PHROGs database.

##### ***B.1.4.*** Microviridae data

A total of 15,507 *Microviridae* genomes were collected from public databases as following: 4,498 genomes were downloaded with their associated metadata from NCBI Datasets on July 8, 2024 with the keyword ‘*Microviridae*’ and the filter ‘complete’ (O’Leary et al., 2024), 130 genomes were downloaded from the PhageScope database on July 31, 2024 with default filters, the taxonomy ‘*Microviridae*’, and sequence quality ‘Complete’ (R. Wang et al., 2024), and 10,879 genomes of the virus family ‘*Microviridae*’ were downloaded from the OpenGenome database (Brixi et al., 2025; Nguyen, Poli, Durrant, et al., 2024). 261 genomes over 10 kb in length or containing characters other than ‘A’, ‘C’, ‘G’, or ‘T’ were removed. The filtered sequences were clustered at 99% identity by MMseqs2 version 13.45111 (Steinegger & Söding, 2017) and a total of 14,466 final sequences were extracted. The relative proportions of available sequencing data for bacteriophage families were analyzed from ‘Complete’ genomes collected from NCBI Virus on May 12, 2025 (Quinones-Olvera, n.d.)(https://github.com/nataquinones/phage_genome_size/). For analyses involving ΦX174 variants, a total of 134 ΦX174 variant sequences were extracted from the raw *Microviridae* data by searching for the keyword ‘phix174’.

##### ***B.1.5.*** Data preprocessing for supervised fine-tuning

*Microviridae* sequences were aligned to the ΦX174 NCBI reference sequence NC_001422.1 using align. align_optimal from the biotite Python package version 0.39.0 (Kunzmann & Hamacher, 2018) and the sequences were prepended with tokens indicating their percent sequence identity to ΦX174. All sequences were prepended with the token “+” indicating *Microviridae*. After the “+” token, sequences with 95–100% identity to ΦX174 were prepended with “∼”, 80–95% with “^”, 70–80% with “#”, 50–70% with “$”, and <50% with “!”. Note that this soft-prompting strategy was not used for the final generation of generated phage candidates; however, the tokens “+∼” were prepended to every prompt before generation. Before finetuning, phage genomes were randomly split into 14,266 training sequences, 100 validation sequences, and 100 test sequences.

##### ***B.1.6.*** Supervised fine-tuning of Evo 1

A full fine-tune of the Evo 1 7B 131K model was conducted for 5,000 iterations across 16 Nvidia H100 GPUs, with a batch size of 64 samples and a context length of 10,240 tokens, corresponding to a global batch size of 655,360 tokens. Each sample corresponded to a single phage genome, including special prompt tokens, where sequences shorter than 10,240 tokens were padded up to the context length and where the training loss was only defined on the sequence, excluding pad tokens. Most of the hyperparameters used during pretraining were retained (Nguyen, Poli, Durrant, et al., 2024) but the initial learning rate was set to 0.00009698 with linear warmup through 5% of the fine-tuning iterations followed by cosine decay through the remaining finetuning iterations to a minimum learning rate of 0.00003.

##### ***B.1.7.*** Supervised fine-tuning of Evo 2

A full fine-tune of the Evo 2 7B 8K model was conducted for 12,000 iterations across 32 Nvidia H100 GPUs, with a batch size of 32 samples and a context length of 10,240 tokens, corresponding to a global batch size of 327,680 tokens. Each sample corresponded to a single phage genome, including special prompt tokens, where sequences shorter than 10,240 tokens were padded up to the context length and where the training loss was only defined on the sequence, excluding pad tokens. Most of the hyperparameters used during pretraining were retained (Brixi et al., 2025) but the initial learning rate was set to 0.00001 with linear warmup through 5% of the fine-tuning iterations followed by cosine decay through the remaining fine-tuning iterations to a minimum learning rate of 0.000001.

##### ***B.1.8.*** Positional entropy analysis

Per-position entropy was calculated as the entropy of the conditional probability 𝑝(𝑥_𝑖_ |𝑥_1_, …, 𝑥_𝑖−1_) predicted by Evo at each position 𝑖 for token 𝑥_𝑖_. The positional entropies of all ΦX174 variants extracted from the *Microviridae* dataset were calculated by Evo 1 7B 131K, Evo 2 7B 8K, and training checkpoints of the Evo 1 SFT and Evo 2 SFT models at 5K steps and 12K steps, respectively, using a custom script (**Data and code availability**). Per-position entropies were averaged across sequences of length 5386 and smoothed with a Savitzky–Golay filter, with a window length of 10 and polynomial order of 3.

##### ***B.1.9.*** Design constraints

Sequences were subject to a variety of nucleotide- and gene-level design constraints to determine their biological feasibility (**Data and code availability**). The design constraints were separated into three categories based on their purpose: quality control, host tropism specificity, and sequence diversification. Quality control constraints included filtering for sequences with only characters A, C, G, T, sequence length of 4–6 kb, 30–65% GC content, DNA homopolymer length of ≤10 nt, and predicted protein hit count ≥7 (**Figure S7A**). Protein sequences were predicted using our custom gene annotation method (see ORF calling and gene annotation). Sequences retained after applying quality control filters were passed to the tropism filter, which kept sequences containing a spike protein with ≥60% protein sequence identity to ΦX174 spike protein, determined by MMseqs2 version 13.45111 (Steinegger & Söding, 2017) with sensitivity = 4.0 (**Figure S7A**). The tropism-filtered sequences were then manually inspected or passed to further diversification filters, including architectural similarity score ≤0.9, average amino acid identity (AAI) ≤95%, synteny break of one gene, and total gene count of 10 or 12 genes (**Figure S7A**). Generated sequences were additionally analyzed, but not filtered with, CheckV version 0.7.0 (Nayfach et al., 2021) with default settings, to determine viral sequence quality. Sequences were visualized by LoVis4u with flags -hl, --set-category-colour, -cA4p2, -alip (Egorov & Atkinson, 2025) to aid manual inspection.

##### ***B.1.10.*** ORF calling and gene annotation

To enable accurate gene-level design constraints, various gene prediction methods were compared using the reference ΦX174 genome NC_001422.1. Prodigal version 2.6.3 was run with the flag the -pmeta (Hyatt et al., 2010). Pyrodigal-gv version 0.3.2 was run with default settings (Camargo et al., 2023). Phanotate version 1.6.5 was run with default settings (McNair et al., 2019). pLannotate was run via the web server (http://plan notate.barricklab.org/) using default parameters (McGuffie & Barrick, 2021). Glimmer was run in Geneious Prime version 2025.1.2 with the following parameters: minimum gene length = 90, maximum overlap length = 1200, start codons = ATG, stop codons = TAG/TGA/TAA, and genetic code = 11 (Delcher et al., 2007). GeneMark annotation was performed using MetaGeneMark version 3.42 via the web server (https://gene mark.bme.gatech.edu/heuristic_gmhmmp.cgi) with default parameters (Besemer & Borodovsky, 2005). These methods failed to predict all 11 genes in the ΦX174 genome, which prompted us to create our own method tailored to ΦX174 (Fig. S4C; Data and code availability). First, genomes were “pseudo-circularized”, by searching for the first stop codon position in each reading frame, identifying the most downstream first stop codon position, extracting the sequence up to that position, and appending it to the end of the genome. Then, all possible ORFs were determined using orfipy (Singh & Wurtele, 2021) with the input start codon ATG, and output stop codons, TAA, TAG, and TGA. Finally, an all-by-all search was performed against the PHROGs database (Terzian et al., 2021) using MMseqs2 version 13.45111 (Steinegger & Söding, 2017) with sensitivity = 4.0. For each hit against the PHROGs database, only the most significant hit (lowest E-value) was kept as the predicted annotation.

##### ***B.1.11.*** Genetic architecture similarity scoring

To evaluate architectural changes in genome sequences (i.e., the arrangement of open reading frames) compared to the reference ΦX174 genome NC_001422.1, we created a genetic architecture scoring algorithm (**Data and code availability**). Since local and global sequence alignment methods based on dynamic programming algorithms are compute-inefficient (Steinegger & Söding, 2017), and percent identity calculated from sequence alignment does not directly inform changes in genetic architecture, we reasoned that a simple dot product scoring method would be a fast way to broadly determine changes in the genetic architectures of sequences. The genes in ΦX174 begin with an ATG start codon and end with a stop codon (TAA, TAG, TGA). Thus, we reasoned that we could quickly compare a generated architecture with the ground truth ΦX174 genome architecture by one-hot encoding start and stop codon positions, creating sparse vectors that represent the boundaries of genes. We created one-hot encoded ‘truth’ vectors of ΦX174: a vector T which is Gaussian blurred (𝜎 = 5) to create fuzzy gene boundaries that tolerate slight positional shifts without over-penalizing near-matches, and a weight vector W that is multiplied by a weight factor equivalent to the total number of gene boundaries. A ‘query’ matrix of the one-hot encoded vectors of all circular permutations of a generated sequence is then scored with NumPy operations (Harris et al., 2020) np.multiply(W,np.max(np.dot(T,Q)))/N, where 𝑁 is the genetic architecture score of the ΦX174. Thus, a score of 1 indicates an exact match to the genetic architecture of ΦX174.

##### ***B.1.12.*** UMAP visualization of genetic architectures

One-hot encoded genetic architecture vectors with the highest genetic architecture similarity score to ΦX174 were computed for genomes in the *Microviridae* training data and stored in an AnnData object (Virshup et al., 2021). Using Scanpy (Wolf et al., 2018), the vectors were Z-score normalized and reduced to 50 principal components by PCA. A 𝑘-nearest neighbor graph (𝑘 = 50) was constructed on the PCA representation, UMAP was applied for non-linear dimensionality reduction, and the resulting UMAP embeddings were visualized.

##### ***B.1.13. Φ***X174 genome prompt analysis

All ΦX174 variants extracted from the *Microviridae* training dataset were aligned with MAFFT (Katoh & Standley, 2013) in Geneious Prime version 2025.1.2 (https://www.geneious.com/). The consensus start sequence was determined by analyzing the multiple sequence alignment by WebLogo (https://weblogo.berkeley.edu/l ogo.cgi) (Crooks et al., 2004). Increasing amounts of nucleotides from the consensus start sequence were used to prompt the Evo 1 7B 131K, Evo 2 7B 1M, Evo 1 SFT, and Evo 2 SFT models with generation temperature = 1.1, top-k = 4, top-p = 1. Prompt lengths spanned 1 to 11 nucleotides, and all prompts were prepended with “+∼” finetuning tokens. Approximately 1,000 sequences of length 6,000 were generated with each prompt.

Percent recall of ΦX174 per prompt was determined as the percent of generated sequences per prompt with a significant alignment against the reference genome ΦX174 NC_001422.1, using MMseqs2 version 13.45111 (Steinegger & Söding, 2017) with sensitivity = 7.5.

##### ***B.1.14.*** Model temperature and prompt sampling sweeps

To determine the optimal parameter combination for phage genome generation with the Evo 1 SFT and Evo 2 SFT models, sampling sweeps were conducted by systematically varying both model temperature and the nucleotide sequence prompt. Temperature sweeps were performed across five configurations: 0.3, 0.5, 0.7, 0.9, and 1.1. Prompt lengths spanned 1 to 11 nucleotides of the ΦX174 consensus start sequence. All prompts were prepended with “+∼” fine-tuning tokens. Approximately 1,000 sequences were sampled per temperature–prompt parameter combination. All sampling runs were performed with top-k = 4 and top-p = 1. The parameter sweeps were evaluated by Shannon diversity (**Figure 2J**), retention rate after filtering (**Figure 2K**), and diversity of architectural similarity score vs. percent spike protein sequence identity (**Figure S6**). Additional sampling sweeps were performed across model checkpoints, fine-tuning tokens, top-k values, and top-p values, but were not used for final evaluation of generation conditions.

##### ***B.1.15.*** Shannon diversity analysis

Shannon entropy was used to quantify sequence diversity (**Data and code availability**) in generated sequences, *Microviridae* genomes, scrambled *Microviridae* genomes, and the ΦX174 reference genome NC_001 422.1. Generated sequences for each temperature and prompt length were filtered with quality control and tropism constraints before being analyzed. For each dataset, sequences were clustered using MMseqs2 version 13.45111 (Steinegger & Söding, 2017) at 99% nucleotide identity, then Shannon entropy was calculated on the resulting distribution of sequences per cluster.

##### ***B.1.16.*** Sequence filtering retention rate analysis

Generated sequences for each temperature and prompt length were filtered with quality control, tropism, and diversification constraints (**Figure S7A**), and the percentage of sequences passing each group of filters (retention rate) was calculated. The maximum retention rate across all temperature and prompt combinations was used to determine model performance. Design constraints were applied to *Microviridae*, scrambled *Microviridae*, and ΦX174 variant sequences as controls.

#### **B.2.** Functional assays of bacteriophage genomes

##### ***B.2.1.*** Biosafety and sterile technique

All experiments involving bacteriophage particles were conducted in a Class II Type A2 biosafety cabinet meeting NSF/ANSI 49, OEM specifications, and ISO 14644-1 standard. Pipettes and a 37 ^◦^C incubator were dedicated for use only with active bacteriophage cultures. Equipment was regularly sterilized with 70% ethanol, 10% bleach, and UV treatment. For all experiments, appropriate PPE was worn and sterile filter pipette tips were used. Any solid or liquid waste produced from experiments was discarded as biohazard waste.

##### ***B.2.2.*** Bacterial and bacteriophage growth media

Unless stated otherwise, bacteria were grown in liquid culture of “LB” consisting of LB Miller broth (Thermo Fisher Scientific #H26676). Bacteriophages were propagated with bacteria in liquid culture of “phage LB” media consisting of 10 g/L tryptone (RPI #T60060), 5 g/L yeast extract (IBI #IB49161), 10 g/L NaCl (Thermo Fisher Scientific #S271), and 2 mM CaCl2 (J.T. Baker #1332). Bacterial colonies and bacteriophage plaques were grown in “phage LB agar” consisting of 7 g/L agar (Carolina #84-2133) in phage LB.

##### ***B.2.3.*** Bacterial strains

Lyophilized *E. coli* C derived from ATCC 13706 (Microbiologics #0747P) was rehydrated following the manufacturer’s protocol in sterile conditions. The stock culture was inoculated in LB, incubated overnight at 37 ^◦^C with 170 rpm agitation, and mixed at a 1:1 volumetric ratio with 50% glycerol (Thermo Fisher Scientific #15514011) and stored at –80 ^◦^C. For preparation of *E. coli* C competent cells, a pipette tip was used to scrape glycerol stock and dipped in 5 mL of LB overnight at 37 ^◦^C with 170 rpm agitation, then 0.5 mL of the overnight culture was used with a Mix & Go *E. coli* Transformation Kit (Zymo Research #T3001) following the manufacturer’s protocol. Competent cell stocks were stored at –80 ^◦^C and thawed only at the time of use. The strains ATCC 25404 (*E. coli* K-12), Stbl3 (*E. coli* K-12/B HB101), JW5856 (*E. coli* K-12 W3110), Stellar (*E. coli* K-12 HST08), RosettaDE3 (*E. coli* B), Mach1 (*E. coli* W), and MDS69 (*E. coli* K-12 with reduced genome) (Karcagi et al., 2016) were a gift from Alex Gao’s lab. Glycerol stocks and competent cell stocks for these strains were prepared the same way.

##### ***B.2.4.*** Bacteriophage genome assembly from ***Φ***X174 RFI DNA

ΦX174 am3 cs70 RFI DNA (NEB #N3021) was PCR-amplified into separate sets of two or three fragments (File S1). The following pairs of primers were used to PCR-amplify fragments for two-fragment Gibson assembly of lysis mutant ΦX174 genomes with the amber (am3) mutation in the lysis gene: SK#327 + SK#324 and SK#323 + SK#328. The following pairs of primers were used to PCR-amplify fragments for two-fragment Gibson assembly of wild-type ΦX174 genomes by reverting the am3 mutation to the wild-type codon: SK#333 + SK#326 and SK#334 + SK#325. The following pairs of primers were used to PCR-amplify fragments for three-fragment Gibson assembly of lysis mutant ΦX174 genomes with the am3 mutation: SK#327 + SK#332 and SK#331 + SK#326 and SK#325 + SK#328. The following pairs of primers were used to PCR-amplify fragments for three-fragment Gibson assembly of wild-type ΦX174 genomes by reverting the am3 mutation to the wild-type codon: SK#333 + SK#322 and SK#321 + SK#318 and SK#317 + SK#334. The fragments were amplified using 1 𝜇L of 1 ng/𝜇L ΦX174 am3 cs70 RFI DNA with 0.5 𝜇L each of 10 uM primers, 10.5 𝜇L of dH2O, and 12.5 𝜇L of 2× CloneAmp HiFi PCR Premix (Takara Bio #639298) with the following settings: 98 ^◦^C for 30 sec, 98 ^◦^C for 10 sec and 62 ^◦^C for 10 sec and 72 ^◦^C for 20 sec repeated 30 times, and 72 ^◦^C for 5 min.

The PCR amplicons were separated in 1% (w/v) UltraPure Agarose (Thermo Fisher Scientific #16500500) in 1× TAE gels stained with 1× SYBR Safe (Thermo Fisher Scientific #S33102) run at 120 V for 30 min, with GeneRuler 1 kb Plus DNA Ladder (Thermo Fisher Scientific #SM1331). PCR products of the correct size were purified by gel extraction using a QIAquick Gel Extraction Kit (Qiagen #28704) following the manufacturer’s protocol. Assembly fragments were mixed with NEBuilder HiFi DNA Assembly Master Mix (NEB #E2621) with the volumes of each fragment calculated on https://nebuildercalculator.neb.com/, and incubated at 50 ^◦^C for 1 hour. Assemblies were stored at –20 ^◦^C until use.

##### ***B.2.5.*** Bacteriophage genome assembly from synthesized DNA

Sequences were split into two Gibson assembly-compatible fragments and synthesized as Gene Fragments by Twist Biosciences, without adapters, dried down and normalized to 250 ng in 96-well plate wells. Optimal split points and overhangs for each Gibson assembly were designed with a custom Python script (**Data and code availability**) that outputs each fragment to be synthesized. The synthesized fragments were resuspended in dH2O to 10, 25, or 50 ng/𝜇L and assembled with the following conditions: each of two fragments per genome were mixed at a 1:1 (v/v) ratio to a total volume of 2 𝜇L. 2 𝜇L of NEBuilder HiFi DNA Assembly Master Mix (NEB #E2621) was added to the mixture on ice to a total reaction volume of 4 𝜇L and the reaction was incubated at 50 ^◦^C for 1 hour. Assemblies were stored at –20 ^◦^C until use. The gene fragments were sealed in their original 96-well plates with foil seals (Bio-Rad #MSF1001) and stored at –20 ^◦^C.

##### ***B.2.6.*** Bacteriophage genome assembly plaque assay

Phage genomes were assembled by Gibson assembly and 4 𝜇L of the assembly was mixed with 100 𝜇L of *E. coli* C competent cells and left on ice for 10 min. The transformation was gently mixed with 0.5 mL of phage LB and added to 7 mL of phage LB agar at a temperature of 42–46 ^◦^C monitored using an Infrared Thermometer (Ketokek #KT600B), then plated immediately on 10 cm Petri dishes. The plates were incubated at 37 ^◦^C for 3 hours, wrapped with a thin strip of Parafilm (Millipore Sigma #HS234526B) to retain moisture, and stored at 4 ^◦^C.

##### ***B.2.7.*** Bacteriophage genome assembly growth assay

Phage genomes were assembled by Gibson assembly and 1 𝜇L of the assembly was mixed with 15 𝜇L of *E. coli* C or *E. coli* K-12 competent cells and left on ice for 10 min. The transformation was gently mixed with 735 𝜇L of phage LB and split into three wells of a flat-bottom 96-well plate (Thermo Fisher Scientific #167008) with 250 𝜇L of per well. Three wells with 250 𝜇L phage LB only were added to the plate to normalize the OD_600_ measurements. The 96-well plate was incubated at 37 ^◦^C with orbital shaking at 1.5 mm amplitude and 360 rpm in a Tecan Spark Multimode Microplate Reader, with OD_600_ measured every 15 min. After 6 hours, the plate was removed from the microplate reader and stored at 4 ^◦^C.

##### ***B.2.8.*** Bacteriophage glycerol stocks

Bacterial debris in phage-clarified cultures were pelleted by centrifugation at 3,000 × g for 10 min at 4 ^◦^C. The supernatant was sterile-filtered through a 0.22 𝜇m cellulose acetate Spin-X Centrifuge Tube Filter (Thermo Fisher Scientific #07-200-385) by centrifugation at 10,000 × g for 3 min at 4 ^◦^C. Glycerol stocks were prepared by mixing the flow-through with 0.22 𝜇m-filtered 50% glycerol (Thermo Fisher Scientific #15514011) at a 1:1 (v/v) ratio and stored at –80 ^◦^C in cryogenic tubes (Thermo Fisher Scientific #11-676-48).

##### ***B.2.9.*** Long-read sequencing of bacteriophage genomes from glycerol stocks

Glycerol stocks prepared from the phage growth assays were individually scraped using sterile pipette tips and dipped into 6 𝜇L of 1× phosphate buffered saline (PBS). The picked phage genomes were amplified using 1 𝜇L of the solution in a PCR reaction with 0.5 𝜇L each of 10 uM primers (File S1), 10.5 𝜇L of dH2O, and 12.5 𝜇L of 2× CloneAmp HiFi PCR Premix (Takara Bio #639298) with the following settings: 98 ^◦^C for 30 sec, 98 ^◦^C for 10 sec and 62 ^◦^C for 10 sec and 72 ^◦^C for 20 sec repeated 30 times, and 72 ^◦^C for 5 min. The PCR amplicons were separated in 1% (w/v) UltraPure Agarose (Thermo Fisher Scientific #16500500) in 1× TAE gels stained with 1× SYBR Safe (Thermo Fisher Scientific #S33102) run at 120 V for 30 min, with GeneRuler 1 kb Plus DNA Ladder (Thermo Fisher Scientific #SM1331). PCR products of the correct size were purified by gel extraction using a QIAquick Gel Extraction Kit (Qiagen #28704) following the manufacturer’s protocol and sequenced using Plasmidsaurus’ Standard Purified Linear/PCR sequencing service. Sequences were aligned by MAFFT v7 (Katoh & Standley, 2013) on Benchling (https://www.benchling.com/).

##### ***B.2.10.*** Bacteriophage propagation and harvesting

A glycerol scrape of *E. coli* C was inoculated in 5 mL of LB and incubated at 37 ^◦^C overnight with 200 rpm agitation. 3 mL of the overnight culture was mixed with 247 mL of phage LB and grown to an OD_600_ of ∼0.4. A scrape of phage glycerol stock was added to the culture and clarified for up to 9 hours. The lysed debris was pelleted by centrifugation at 4,000 × g for 10 min at 4 ^◦^C and sterile-filtered through a 0.22 𝜇m pore size PES membrane filter (Millipore Sigma #S2GPU05RE). The phages Evo-Φ46, Evo-Φ111, and Evo-Φ114 were double propagated due to low titer, by adding 3 mL of the first harvest to 297 mL of phage LB and incubating for 9 hours. The phages Evo-Φ36 and Evo-Φ108 were similarly propagated but with a third propagation due to low titer.

##### ***B.2.11.*** Bacteriophage titering

A glycerol scrape of *E. coli* C was inoculated in 5 mL of LB and incubated at 37 ^◦^C overnight with 200 rpm agitation. 50 𝜇L of the overnight culture was mixed with 5 mL of phage LB and incubated at 37 ^◦^C with 200 rpm agitation. The culture was grown to an OD_600_ of 0.3–0.4 and 675 𝜇L of the culture was added to 22.5 mL of phage LB agar at a temperature of 42–46 ^◦^C. The temperature was monitored using an Infrared Thermometer (Ketokek #KT600B). The mixture was poured in a 15 cm Petri dish and allowed to solidify for up to 5 min. 2 𝜇L of 10-fold serial dilutions of phage in phage LB from 100 to 10-10 was spotted on the agar and fully dried for up to 15 min. An additional 2 𝜇L spot of phage LB only was also spotted as a negative control. The plates were inverted and incubated at 37 ^◦^C for 3 hours, imaged, wrapped with a thin strip of Parafilm (Millipore Sigma #HS234526B) to retain moisture, and stored at 4 ^◦^C. Images were taken on an iPhone with a custom backlight setup and converted to black and white by decreasing the saturation to 0. Individual visible plaques were counted at the highest dilution where they were present across all three replicates, and the titer (plaque forming units (PFU) / mL) per phage was calculated as ((average plaque count / spot volume (mL)) × dilution factor).

##### ***B.2.12.*** Host tropism assay

For each *E. coli* strain, a glycerol scrape was inoculated in 5 mL of LB and incubated at 37 ^◦^C overnight with 200 rpm agitation. 250 𝜇L of the overnight culture was mixed with 25 mL of phage LB and incubated at 37 ^◦^C with 200 rpm agitation. The culture was grown to an OD_600_ of 0.4–0.6 and 200 𝜇L per well was plated in a flat-bottom 96-well plate (Thermo Fisher Scientific #167008). 50 𝜇L of phage stock at a concentration of ∼105 PFU/mL was then added to each well. As negative controls, 50 𝜇L of phage LB was added to each well instead of phage. Three wells with 250 𝜇L phage LB only were added to the plate to normalize the OD_600_ measurements. The 96-well plate was incubated at 37 ^◦^C with orbital shaking at 1.5 mm amplitude and 360 rpm in a Tecan Spark Multimode Microplate Reader, with OD_600_ measured every 15 min. After ∼12 hours, the plate was removed from the microplate reader and stored at 4 ^◦^C.

#### **B.3.** Phylogenetic analysis of bacteriophages

##### ***B.3.1.*** Natural reference genomes

Unless otherwise noted, the wild-type ΦX174 reference genome used for all phylogenetic analyses was sourced from NCBI accession NC_001422.1, and the wild-type G4 reference genome used for all phylogenetic analyses was sourced from NCBI accession NC_001420.2.

##### ***B.3.2.*** Synteny analysis

Synteny was visualized by LoVis4u with flags -hl, --set-category-colour, -cA4p2, -alip, (Egorov & Atkinson, 2025) using GFF3 files for each phage genome created with a custom script (**Data and code availability**). Pairwise synteny between each phage genome was determined in the order presented (**Figure 4A**) and consolidated into a single synteny plot. Since our gene annotation method only partially predicted gene A*, it was omitted from synteny visualization. Synonymous, nonsynonymous, and noncoding mutations were determined by aligning each generated genome against the ΦX174 reference genome with MAFFT (Katoh & Standley, 2013) in Geneious Prime version 2025.1.2 (https://www.geneious.com/), setting ΦX174 as the reference sequence, finding variations/SNVs with Inside\&OutsideCDS and genetic code set as Bacter ial, then colored by ProteinEffect. If synonymous and nonsynonymous mutations overlapped due to overlapping genes, the nonsynonymous mutation was visualized as the top layer. Genes sharing synteny with ΦX174 were determined using a custom script (**Data and code availability**) analyzing the pairwise protein identity matrix calculated by LoVis4u. For simply visualizing SNVs relative to ΦX174, ΦX174 was set as the reference sequence and the default mutation highlighting in Geneious Prime was exported and overlaid on the synteny visualization.

##### ***B.3.3.*** Whole-genome alignment

Unless otherwise noted, whole-genome alignments were performed with MAFFT (Katoh & Standley, 2013) in Geneious Prime version 2025.1.2 (https://www.geneious.com/). To determine percent genome identity of generated phage genome candidates compared to ΦX174 and sequences in the *Microviridae* training data, genomes were aligned by nucleotide BLAST (blastn, E-value = 0.5, default settings) (Camacho et al., 2009) in Geneious Prime, and percent genome identity was calculated as (percent identity × percent query coverage). Percent genome identity of sequences in the *Microviridae* training data compared to ΦX174 were determined by MMseqs2 version 13.45111 (Steinegger & Söding, 2017) with default settings.

##### ***B.3.4.*** Mutational count to nearest natural genome analysis

Generated phage genome candidates were aligned to sequences in the *Microviridae* training data by nucleotide BLAST (blastn, E-value = 0.5, default settings) (Camacho et al., 2009) in Geneious Prime version 2025.1.2 (https://www.geneious.com/). Percent genome identity of each generated sequence to its nearest natural sequence was calculated as (percent identity × percent query coverage). The number of novel nucleotide mutations in each generated genome was estimated as ((1 (percent genome identity / 100)) × generated genome length).

##### ***B.3.5.*** Cumulative genome attribution analysis

To determine if mutations in the generated genomes could be attributed to existing mutations in natural genomes, the top 1,000 alignments (E value < 1.0) for each generated genome were determined using nucleotide BLAST (blastn, E-value = 0.05, default settings) (Camacho et al., 2009). Base pair-level attributions followed a greedy approach, where exact nucleotide matches were first assigned to the highest-scoring BLAST hit, then remaining unassigned positions were iteratively assigned to lower-scoring hits in descending order until complete assignment or no additional matches were possible.

##### ***B.3.6.*** Phylogenetic tree construction

A multiple sequence alignment of representative *Microviridae* and generated phage genomes was constructed with MAFFT (Katoh & Standley, 2013) in Geneious Prime version 2025.1.2 (https://www.geneious.com/). The resulting MSA was used to build a Neighbor-Joining tree with the Geneious Tree Builder tool with a Jukes-Cantor genetic distance model and no outgroup. The following natural reference genomes collected from NCBI were used for phylogenetic trees: NC_001422.1 (ΦX174), KY653237.1 (alpha-𝛼), DQ079890.1 (NC41), DQ079885.1 (NC5), DQ079891.1 (NC51), and AF274751.1 (S13), NC_001420.2 (G4), NC_012868.1 (St-1), NC_001730.1 (ΦK), NC_007821.1 (WA13), NC_001330.1 (𝛼3).

##### ***B.3.7.*** Mutational hotspot analysis

Nucleotide sequences corresponding to annotated genes, promoters, and terminators from wild-type ΦX174 (Logel & Jaschke, 2020) were aligned against all generated genomes using nucleotide BLAST version 2.16.0+ (blastn-short, ungapped, word_size = 4, E-value = 0.2) (Camacho et al., 2009). For each alignment, total mutations were defined as the sum of reported mismatches and unaligned query residues, with the overall mutation rate calculated as the number of total mutations divided by the length of the query sequence. For each gene, promoter, and terminator, the mean mutation rate was calculated by averaging individual mutation rates across all viable generated genomes.

#### **B.4.** Structural analysis of bacteriophages

##### ***B.4.1.*** Protein structure prediction

F (capsid), G (spike) and J (DNA packaging) proteins were co-folded by AlphaFold 3 (Abramson et al., 2024) via the online web server (https://alphafoldserver.com/). PAE was visualized with https://thecodingbiologist.com/tools/pae.html and predicted structures were visualized with UCSF ChimeraX version 1.7.1 (Pettersen et al., 2021).

##### ***B.4.2.*** Bacteriophage purification for cryo-EM

*E. coli* C was grown in 300 mL of phage LB at 37 ^◦^C with 200 rpm agitation until reaching OD_600_ 0.4–0.6, at which they were inoculated with 3 mL of phage for 3.5–8 hours. Chloroform was added to cultures to a final dilution of 1% and incubated for 15 min at 25 ^◦^C with 200 rpm agitation for complete lysis. Lysed cultures were pelleted by centrifugation (3,000 × g, 10 min, 4 ^◦^C), and sterile-filtered through a 0.22 𝜇m pore size PES membrane filter (Millipore Sigma #S2GPU05RE). Filtered lysate was supplemented with polyethylene glycol 8,000, pH 7.4 (8%), 100 mM NaCl and incubated with end-over-end rotation at 4 ^◦^C for 1 hour, before incubation at 4 ^◦^C for 2–16 hours without rotation. The phage pellet was obtained by centrifugation (11,000 × g, 20 min, 4 ^◦^C) and resuspended in 4 mL PBS pH 7.4, 10 mM MgCl2, to which 50 U/mL Benzonase Nuclease (Teknova #E1014) was subsequently added. The mixture was incubated for 45 min at 37 ^◦^C, with gentle mixing every 15 min. The phage suspension was carefully placed on top of a manually constructed discontinuous iodixanol (Teknova #21449, #21443, #21431, #21425) density gradient (60%, 40%, 25%, 15%), and subjected to ultracentrifugation (200,000 × g, 2–4 hours, 4 ^◦^C). Visible phage bands (at the 60% / 40% fraction boundary) were carefully extracted and buffer exchanged by four rounds of centrifugation (15,000 × g, 10 min, 4 ^◦^C) in 100 kDa molecular weight cut-off Amicon centrifugal filters (Millipore #UFC510024) followed by resuspension in 20 mM Tris pH 7.4, 100 mM NaCl.

##### ***B.4.3.*** Polyacrylamide gel electrophoresis (PAGE)

SDS-PAGE was performed using 4–20% SurePAGE precast gels (GenScript #M00657) in XCell SureLock Mini-cell gel chambers (Thermo Fisher Scientific #EI0001). Before loading onto gels, samples were mixed with 4× Laemmli buffer (62.5 mM Tris HCl, pH 6.8, containing 2% (w/v) SDS, 10% (v/v) glycerol, and 0.002% (w/v) bromophenol blue) and boiled at 99 ^◦^C for 3 min. Gels were run at 200 V in 1× MES SDS Running Buffer (Genscript #M00677) alongside SeeBlue Plus2 Pre-stained Protein Standard ladder (Invitrogen #LC5925) or PageRuler Plus Prestained Protein ladder (Thermo Fisher Scientific #26619). Gels were stained with Instant-Blue Coomassie (AbCam #ab119211) for up to 1 hour with gentle shaking and de-stained for at least 1 hour with Milli-Q water prior to imaging on a GelDoc Go imager (Bio-Rad). ImageLab version 6.1.0 (Bio-Rad) was used for analysis.

##### ***B.4.4.*** Grid preparation for cryo-EM

Phage samples were purified as described above and stored at 4 ^◦^C prior to cryo-EM grid preparation. 3.2 𝜇L of the purified phage was applied to R1.2/1.3, 300 mesh carbon Cu grids (QUANTIFOIL #Q3100CR1.3) which were glow-discharged using a PELCO easiGlow system with 10 mA negative current for a total period of 90 s at 0.26 mBar with a 10 s hold period. The grids were plunge-frozen in liquid ethane using a Vitrobot Mark IV (Thermo Fisher Scientific) maintained at 100% humidity and 8 ^◦^C with a blot time of 3 s and a blot force of 3.5. The grids were stored in liquid nitrogen.

##### ***B.4.5.*** Cryo-EM data acquisition

Grids of ΦX174 and Evo-Φ36 were initially screened on a Glacios electron microscope (Thermo Fisher Scientific). One high-quality grid for each phage was selected for data collection and imaged on a Glacios electron microscope (Thermo Fisher Scientific) operated at 200 kV and equipped with a Falcon 4i direct electron detector. Movies were automatically collected with EPU software (Thermo Fisher Scientific) at 150,000× magnification magnification, corresponding to a real pixel size of 0.923 Å at the specimen level. For ΦX174, 3,372 movies were collected with a defocus range of -1.5 to -3.0 microns. Each movie was recorded over 3.52 s with a total accumulated dose of 41.15 𝑒^−^/Å^2^, fractionated into 40 frames (∼1.03 𝑒^−^/Å^2^ per frame). For Evo-Φ36, 4,796 movies were collected with a defocus range of -1.5 to -2.5 microns. Each movie was recorded over 3.79 s with a total accumulated dose of 50.00 𝑒^−^/Å^2^, fractionated into 43 frames (∼1.16 𝑒^−^/Å^2^ per frame).

##### ***B.4.6.*** Cryo-EM data processing

Cryo-EM data were processed using cryoSPARC version 4.6.2 (Punjani et al., 2017). Movies were motion-corrected and CTF parameters estimated using the patch motion correction and patch-based CTF estimation jobs. Initial particles were picked on denoised micrographs by blob picking using a circular blob with particle diameter between 250 Å and 325 Å. Picked particles were extracted with a box size of 512 pixels and classified in 2D to select representative 2D classes for use as templates. Resulting particles from template-guided particle picking were extracted with a box size of 512 pixels and subjected to 2–3 rounds of 2D classification, yielding 24,999 particles for ΦX174 and 30,228 particles for Evo-Φ36. Multiple rounds of three-dimensional (3D) reconstruction and refinement (uniform and non-uniform with icosahedral symmetry imposed, as well as global and local CTF refinement) were performed without further particle sorting, with one round of reference-based motion correction, with 24,547 particles and 29,708 particles used in the final refinements for ΦX174 and Evo-Φ36 respectively. This resulted in final maps for ΦX174 and Evo-Φ36 with resolutions of 2.76 Åand 2.90 Å, respectively, as determined by their gold standard Fourier shell correlation with a threshold of 0.143 (Rosenthal & Henderson, 2003). Further processing details are provided in **Figure S17**. Final maps for model building were sharpened using a postprocessing job in RELION 5.0 with masks constructed in RELION using a low-pass filter of 20 Å, 1 pixel extension, and 8 pixel soft-edge with a binarization threshold of 0.015 and 0.01 for ΦX174 and Evo-Φ36 respectively (Kimanius et al., 2021).

##### ***B.4.7.*** Cryo-EM model building

AlphaFold3 models of the F, G, and J proteins co-folded for either ΦX174 or Evo-Φ36 were docked into their respective sharpened cryo-EM densities in UCSF ChimeraX before manual adjustment in Coot version 0.9.8.96 (Emsley & Cowtan, 2004). Interactive molecular dynamics-based refinement was performed in ISOLDE version 1.9 (Croll, 2018) to correct local geometry and improve fit to density. Final refinement was carried out in Phenix version 1.21.2 (Liebschner et al., 2019) against the cryo-EM density maps, applying appropriate geometry and secondary-structure restraints. Structures were visualized with UCSF ChimeraX version 1.7.1 or 1.9.1 (Pettersen et al., 2021).

#### **B.5.** Fitness assays of bacteriophage genomes

##### ***B.5.1.*** Bacteriophage competition assay

A glycerol scrape of *E. coli* C was inoculated in 100 mL of LB and incubated at 37 ^◦^C overnight with 200 rpm agitation. 2 𝜇L of the overnight culture was mixed with 198 𝜇L of phage LB in a flat-bottom 96-well plate (Thermo Fisher Scientific #167008) wells and incubated at 37 ^◦^C with orbital shaking at 1.5 mm amplitude and 360 rpm in a Tecan Spark Multimode Microplate Reader. Three wells with 250 𝜇L phage LB only were added to the plate to normalize the OD_600_ measurements. Once the cultures reached OD_600_ ∼0.4, 50 𝜇L of a phage mixture consisting of ∼1×105 PFU/mL per phage of all 16 generated phages and ΦX174 was added to each well. The plate was again incubated in the microplate reader with the same conditions, with OD_600_ measured every 10 min. Upon the initial infection, 10 𝜇L was extracted from each culture, then 10 𝜇L was extracted every 30 min for 3 hours, then every 60 min for 3 hours, for a total of 6 hours. Each extraction was immediately boiled in a thermal cycler at 98 ^◦^C for 45 s and stored in –20 ^◦^C. 2 𝜇L of the lysate at each time point was used in a PCR reaction with 1 𝜇L each of 10 uM primers SK#324 and SK#359, 21 𝜇L of dH2O, and 25 𝜇L of 2× CloneAmp HiFi PCR Premix (Takara Bio #639298) with the following settings: 98 ^◦^C for 30 sec, 98 ^◦^C for 10 sec and 62 ^◦^C for 10 sec and 72 ^◦^C for 40 sec repeated 25 times, and 72 ^◦^C for 5 min. The PCR amplicons were separated in 1.5% (w/v) UltraPure Agarose (Thermo Fisher Scientific #16500500) in 1× TAE gels stained with 1× SYBR Safe (Thermo Fisher Scientific #S33102) run at 120 V for 30 min, with GeneRuler 1 kb Plus DNA Ladder (Thermo Fisher Scientific #SM1331). PCR products of the correct size were purified by gel extraction using a QIAquick Gel Extraction Kit (Qiagen #28704) following the manufacturer’s protocol and sequenced using Plasmidsaurus’ Standard PCR Premium sequencing service.

##### ***B.5.2.*** Bacteriophage competition sequencing analysis

The raw sequencing reads from each competition time point were analyzed using a custom script (**Data and code availability**). Sequencing reads with a MAPQ score of <20, an alignment length of <70%, and a percent identity of <90% against their top alignment of the generated phage genomes and ΦX174 were filtered out from the analysis. The top alignment hit was considered the phage identity of a given sequencing read. The fold change for a given phage genome at each time point tn from the time point before it tn-1 was calculated as the read count at tn divided by the read count at tn-1. The cumulative fold change for a given phage genome at time point tn was calculated as the sum of the fold change of all previous time points since the initial infection at t0.

##### ***B.5.3.*** Bacterial growth rate assay

A glycerol scrape of *E. coli* C was inoculated in 5 mL of LB and incubated at 37 ^◦^C overnight with 200 rpm agitation. 250 𝜇L of the overnight culture was mixed with 25 mL of phage LB and incubated at 37 ^◦^C with 200 rpm agitation. The culture was grown to an OD_600_ of ∼0.4 and 200 𝜇L per well was plated in a flat-bottom 96-well plate (Thermo Fisher Scientific #167008). 50 𝜇L of each phage stock at a concentration of ∼105 PFU/mL was then added to each well. As negative controls, 50 𝜇L of phage LB was added to each well instead of phage.

Three wells with 250 𝜇L phage LB only were added to the plate to normalize the OD_600_ measurements. The 96-well plate was incubated at 37 ^◦^C with orbital shaking at 1.5 mm amplitude and 360 rpm in a Tecan Spark

Multimode Microplate Reader, with OD_600_ measured every 15 min. After ∼12 hours, the plate was removed from the microplate reader and stored at 4 ^◦^C.

##### ***B.5.4.*** Bacterial growth rate determination

Bacterial growth rates were derived from OD_600_ measurements collected at fixed time intervals. For each replicate, growth rate was computed as the numerical derivative of OD_600_ with respect to time using NumPy’s operation np.gradient in Python (Harris et al., 2020), yielding instantaneous rates expressed as ΔOD_600_ per minute. Replicate trajectories were then aggregated to calculate summary statistics (mean, standard deviation, and minimum growth rates), which were used for downstream comparative analyses across phage treatments.

##### ***B.5.5.*** Statistical analysis

Statistical analyses were performed in Python using the statsmodels package (Seabold & Perktold, 2010). For each assay metric (signed area under the curve, minimum OD_600_ after peak, time to post-peak minimum, and minimum growth rate), replicate-level values were first cleaned by removing missing values. A type II one-way analysis of variance (ANOVA) was used to test for overall differences across phage groups, with statistical significance set at 𝛼 = 0.05. Where the ANOVA indicated significance, Tukey’s honestly significant difference (HSD) test was applied for all pairwise comparisons to identify specific differences between groups. Replicate distributions were visualized alongside group means and standard deviations.

##### ***B.5.6.*** Bacteriophage resistance assay

ΦX174 was assembled by Gibson assembly and 1 𝜇L of the assembly was mixed with 15 𝜇L of *E. coli* C competent cells and left on ice for 10 min. The transformation was gently mixed with 735 𝜇L of phage LB and split into three wells of a flat-bottom 96-well plate (Thermo Fisher Scientific #167008) with 250 𝜇L per well. 15 𝜇L of non-transformed competent cells were mixed with 735 𝜇L of phage LB and plated in another three wells. Three wells with 250 𝜇L phage LB only were added to the plate to normalize the OD_600_ measurements. The 96-well plate was incubated at 37 ^◦^C with orbital shaking at 1.5 mm amplitude and 360 rpm in a Tecan

Spark Multimode Microplate Reader, with OD_600_ measured every 15 min. After 24 hours, the transformed cultures reached stable resistance against ΦX174, and the non-transformed, ΦX174-susceptible cultures also reached equilibrium growth. 250 𝜇L of each culture was mixed with 50% glycerol (Thermo Fisher Scientific #15514011) at a 1:1 (v/v) ratio and stored at –80 ^◦^C. To isolate single colonies from each strain, the glycerol stocks were individually streaked out on phage LB agar plates and incubated at 37 ^◦^C overnight. Colonies were picked, dipped in 250 𝜇L of phage LB in a 96-well plate, and incubated at 37 ^◦^C overnight with 200 rpm agitation. 50 𝜇L of each culture was mixed with 50% glycerol at a 1:1 (v/v) ratio and stored at –80 ^◦^C for final stocks of the resistant and susceptible *E. coli* C strains. Each strain’s genome was sequenced using Plasmidsaurus’ Standard Bacteria Genome with Extraction service.

To test the ability of ΦX174 and generated phages to inhibit growth of the resistant and susceptible strains, we devised a bacteriophage counter-resistance evolution assay similar to a previously described protocol (Romeyer Dherbey et al., 2023). The following mixtures of phages were prepared: ΦX174 diluted to a concentration of ∼17 × 105 PFU/mL in phage LB (“ΦX174-only cocktail”), and all 16 generated phages and ΦX174 diluted together in phage LB each at a concentration of ∼1 × 105 PFU/mL (“generated phage cocktail”). A scrape of each glycerol stock of the resistant and susceptible strains were inoculated in 250 𝜇L phage LB and incubated at 37 ^◦^C with 200 rpm agitation overnight. For each resistant strain tested, three different conditions were prepared in a 96-well “seed” plate: a “mixed well” consisting of 1 𝜇L of overnight resistant culture with 1 𝜇L of overnight susceptible culture in 198 𝜇L of phage LB (to facilitate evolution of the phages in susceptible cells in the presence of resistant cells), a resistant-only well consisting of 2 𝜇L of overnight resistant culture in 198 𝜇L of phage LB, and a susceptible-only well consisting of 2 𝜇L of overnight susceptible culture in 198 𝜇L of phage LB. The plate was incubated at 37 ^◦^C with orbital shaking at 1.5 mm amplitude and 360 rpm in a Tecan Spark Multimode Microplate Reader, with OD_600_ measured every 15 min until the cultures reached OD_600_ ∼0.4. 50 𝜇L of the ΦX174-only cocktail and the generated phage cocktail were added to the wells as initial infections, and three wells with 250 𝜇L phage LB only were added to the plate to normalize the OD_600_ measurements. The plate was again incubated in the microplate reader the same way, for approximately 3 hours. Resistant-only wells were observed for growth inhibition, and in the meantime, another seed plate was prepared. Each culture in the phage-infected plate was transferred into a 96-well plate (Thermo Fisher Scientific #N8010560) and the cells were pelleted at 3,000 × g for 10 min at 4 ^◦^C. The supernatants were collected and 50 𝜇L of the supernatant from each mixed well was passaged into their corresponding resistant conditions in the new seed plate. Passages were repeated five times. At each passage, any resistant-only culture with inhibited growth due to a successfully counter-evolved phage cocktail was harvested: the cells were pelleted at 3,000 × g for 10 min at 4 ^◦^C, and the supernatant was sterile-filtered through a 0.22 𝜇m cellulose acetate Spin-X Centrifuge Tube Filter (Thermo Fisher Scientific #07-200-385) by centrifugation at 10,000 × g for 3 min at 4 ^◦^C. The harvested phage cocktails were stored at 4 ^◦^C.

##### ***B.5.7.*** Bacteriophage cocktail sequencing

A glycerol scrape of resistant *E. coli* C corresponding to each evolved phage cocktail was inoculated in 5 mL of LB and incubated at 37 ^◦^C overnight with 200 rpm agitation. 50 𝜇L of the overnight culture was mixed with 5 mL of phage LB and incubated at 37 ^◦^C with 200 rpm agitation. The culture was grown to an OD_600_ of ∼0.4 and 300 𝜇L of the culture was added to 7 mL phage LB agar at a temperature of 42–46 ^◦^C monitored using an Infrared Thermometer (Ketokek #KT600B). To sequence individual phages in the evolved phage cocktails, a pipette tip was dipped in the cocktails and scraped on the surface of the agar plate and incubated at 37 ^◦^C for 3 hours. Eight individual plaques per plate were randomly picked and dipped in 6 𝜇L of phage LB. 2 𝜇L of the phage LB was used per PCR reaction with 1 𝜇L each of 10 uM primers (File S1), 21 𝜇L of dH2O, and 25 𝜇L of 2× CloneAmp HiFi PCR Premix (Takara Bio #639298) with the following settings: 98 ^◦^C for 30 sec, 98 ^◦^C for 10 sec and 62 ^◦^C for 10 sec and 72 ^◦^C for 40 sec repeated 25 times, and 72 ^◦^C for 5 min. The PCR amplicons were separated in 1.5% (w/v) UltraPure Agarose (Thermo Fisher Scientific #16500500) in 1× TAE gels stained with 1× SYBR Safe (Thermo Fisher Scientific #S33102) run at 120 V for 30 min, with GeneRuler 1 kb Plus DNA Ladder (Thermo Fisher Scientific #SM1331). PCR products of the correct size were purified by gel extraction using a QIAquick Gel Extraction Kit (Qiagen #28704) following the manufacturer’s protocol and sequenced using Plasmidsaurus’ Standard Purified Linear/PCR sequencing service. The sequencing fragments were aligned against all generated phage genomes and ΦX174 by MAFFT (Katoh & Standley, 2013) with default settings in Geneious Prime version 2025.1.2 (https://www.geneious.com/) and analyzed for regions of homology. Evo-ΦR1 was chosen as the sequence in which eight out of eight plaques were identical from the cocktail evolved against resistant strain 1, Evo-ΦR2 was chosen as the sequence in which six out of eight plaques were identical from the cocktail against resistant strain 2.

##### ***B.5.8.*** Whole-genome sequencing analysis of ***Φ***X174-resistant E. coli strains

To identify mutational differences potentially conferring resistance amongst ΦX174-resistant *E. coli* C strains, protein FASTA files generated by Bakta (Schwengers et al., 2021), provided by Plasmidsaurus’ Standard Bacteria Genome with Extraction service, for resistant and susceptible *E. coli* C populations were compared. Briefly, each FASTA file was parsed to extract standardized protein descriptors and sequences. For all proteins with identical descriptors, corresponding amino acid sequences were initially compared using simple string equality, with identical sequences being classified as exact matches between resistant and susceptible strains. Non-identical protein sequences were subsequently aligned using MAFFT version 7.525 (Katoh & Standley, 2013) to identify specific amino acid-level differences. Any remaining proteins with non-matching descriptors between the resistant and susceptible strains were then aligned against each other using MAFFT to account for potential large truncations, insertions, or deletions. Differences were then compiled and visualized using Matplotlib.

---

**Figure S1.**
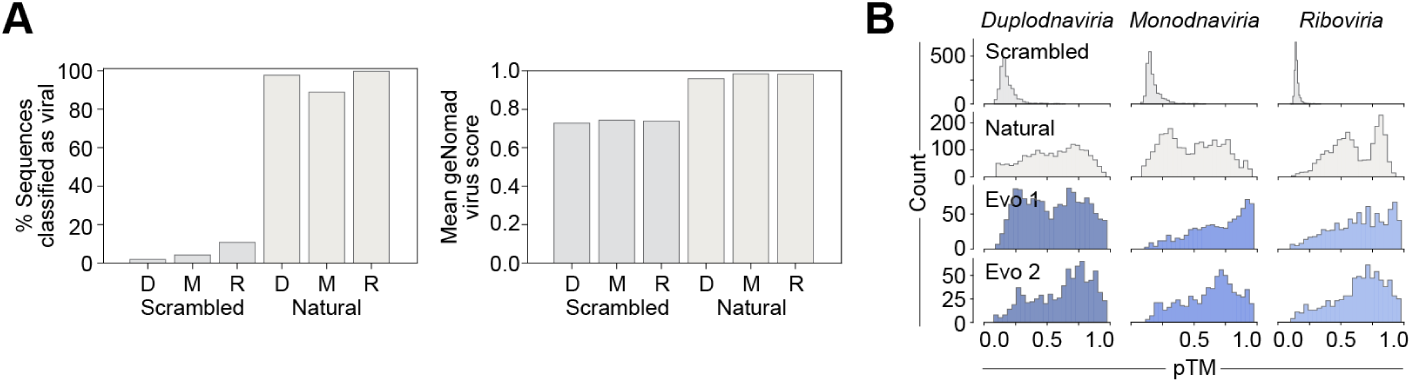
Supplementary data for phage realm sequence generation. **(A)** Percent of natural and scrambled natural sequences classified as viral by geNomad (left), and their mean geNomad virus scores (right). D, *Duplodnaviria*; M, *Monodnaviria*; R, *Riboviria*. geNomad effectively discriminates real natural sequences from scrambled ones. **(B)** ESMFold-predicted protein structures from generated sequences have mean predicted template modeling (pTM) scores similar to natural proteins, substantially higher than scrambled natural sequence controls.

**Figure S2.**
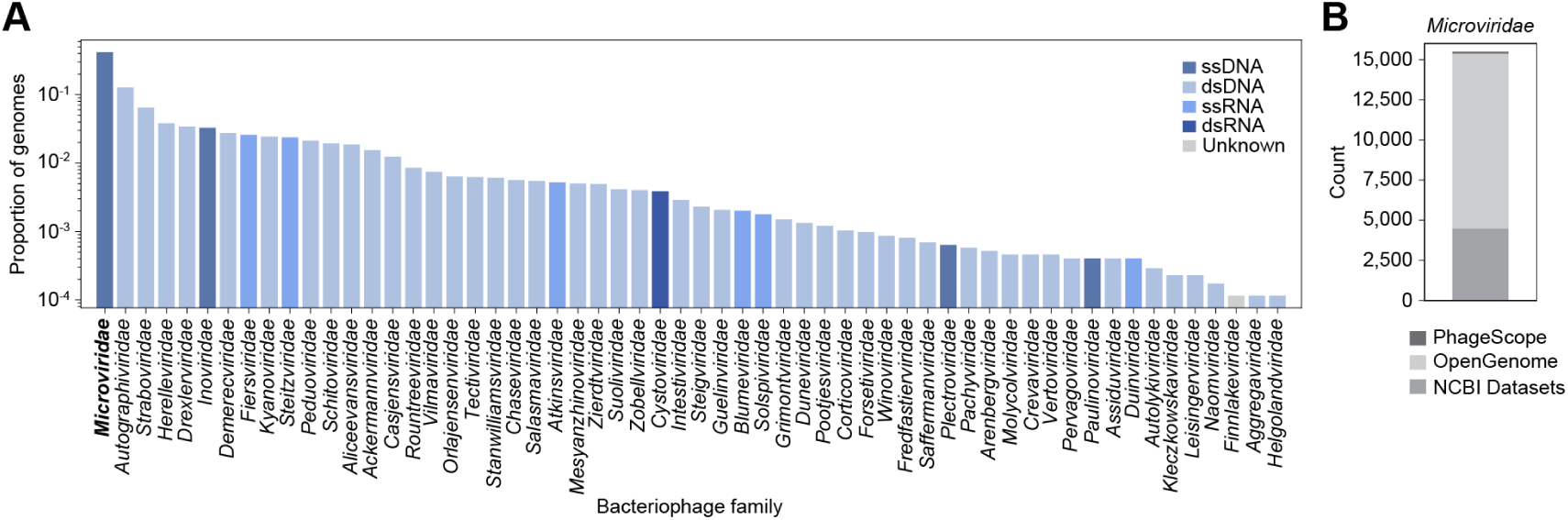
***Microviridae* sequence data. (A)** Relative proportions of complete phage genomes on NCBI Virus as of May 12, 2025. **(B)** *Microviridae* data for fine-tuning collected from PhageScope, OpenGenome, and NCBI Datasets.

**Figure S3.**
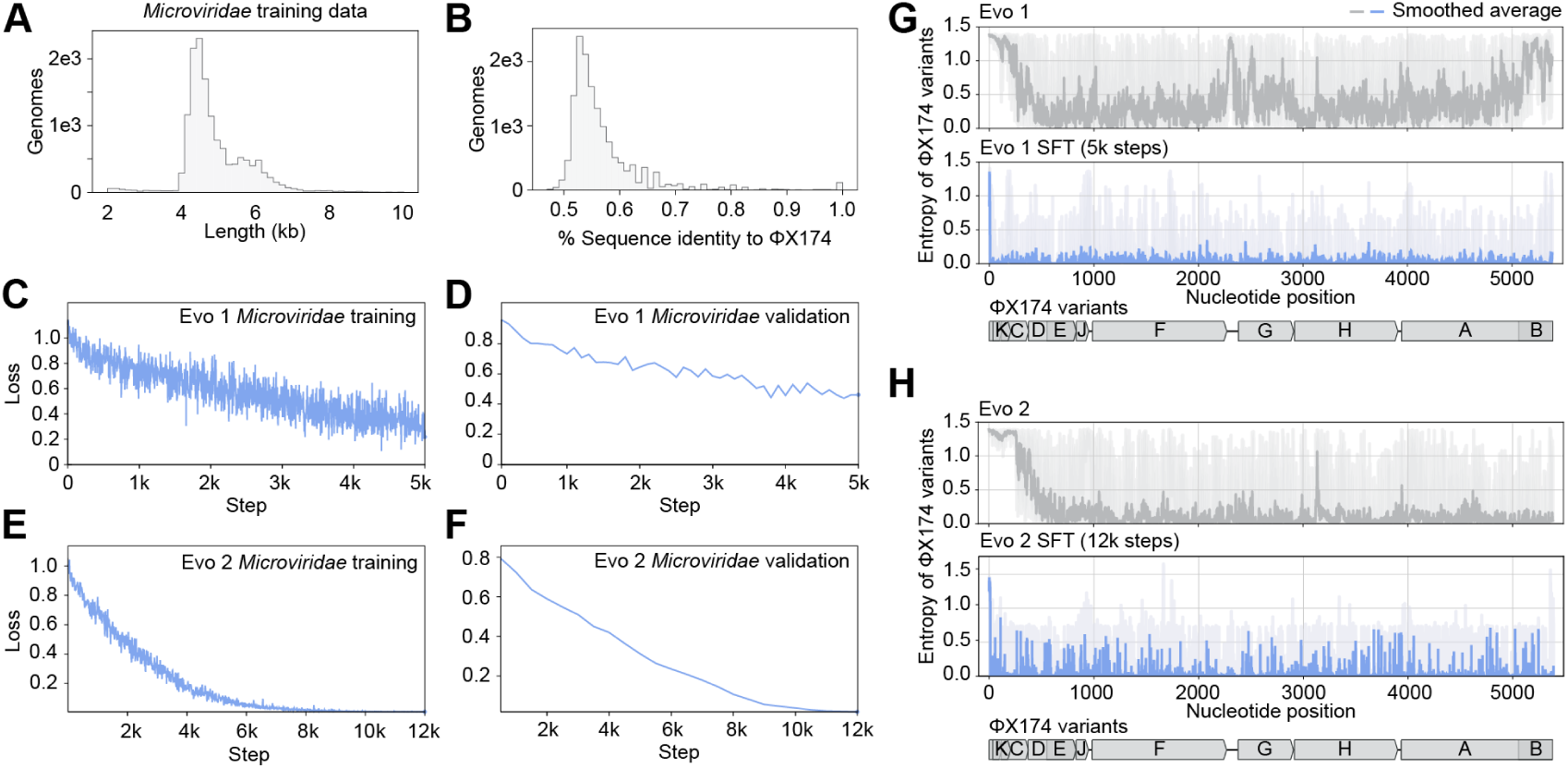
Fine-tuning Evo 1 and Evo 2 on *Microviridae* genomes. **(A)** Lengths of genomic sequences in the *Microviridae* training data. kb, kilobase. **(B)** Percent sequence identity of genomes in the *Microviridae* data aligned to ΦX174. **(C–F)** Training and validation loss curves for Evo 1 fine-tuning **(C–D)** and Evo 2 fine-tuning **(E–F)**. **(G–H)** Per-position entropy of ΦX174 variant sequences calculated by Evo 1 7B 131K and Evo 1 SFT at 5k training steps **(G)**, and Evo 2 7B 8K and Evo 2 SFT at 12k training steps **(H)**. Dark gray and dark blue, smoothed average of positional entropies; light gray and light blue, range of position entropies. Schematic of the ΦX174 genome is shown below.

**Figure S4.**
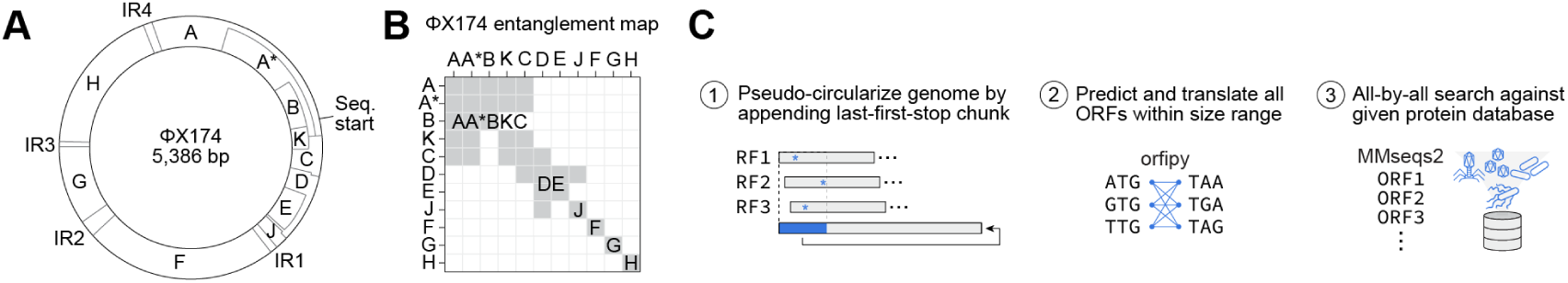
Method for predicting genes in. Φ**X174-like sequences. (A)** Circular schematic of the ΦX174 genome. IR, intergenic region. **(B)** Pairwise matrix indicating overlapping genes (gray) in the ΦX174 genome. **(C)** Steps of our gene annotation method. First, sequences are “pseudo-circularized” by searching for the first stop codon in each reading frame (RF), identifying the most downstream first stop codon position, extracting the sequence up to the position, and appending it to the end of the genome. Then, all possible open reading frames (ORFs) are determined with input start and stop codons using orfipy (Singh & Wurtele, 2021). Finally, an all-by-all search of the ORFs against an input protein database using MMseqs2 (Steinegger & Söding, 2017) is performed.

**Figure S5.**
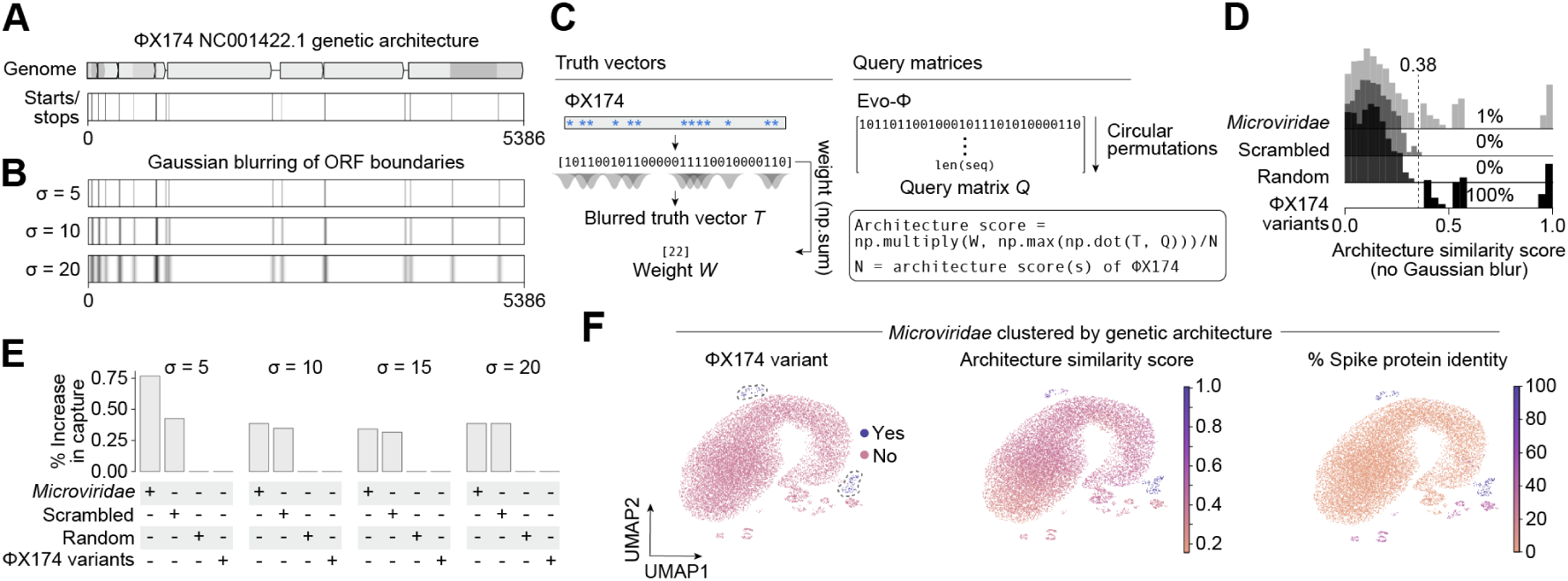
Scoring architectural similarity of genomes. **(A)** Top: Schematic of ΦX174 NC_001422.1 genome architecture. Bottom: Visualization of one-hot encoding of the known start and stop codon boundaries in the ΦX174 genome. **(B)** Applying Gaussian blurs to the one-hot encoding produces smoothed open reading frame (ORF) boundary profiles, making the similarity metric less sensitive to exact start/stop positions. **(C)** Architectural similarity scoring process (**Methods**). **(D)** Distribution of architecture similarity scores normalized to ΦX174 NC_001422.1 without Gaussian blurring show that ΦX174 variants score highly (close to 1), while natural *Microviridae*, scrambled *Microviridae*, and random sequences have low scores. A score >0.38 reliably delineates ΦX174-like sequences. **(E)** Increasing the Gaussian blur parameter 𝜎 expands the capture of sequences above the 0.38 threshold. At 𝜎 = 5, the method balances sensitivity (capturing more ΦX174-like variants) with specificity (limiting scrambled or random sequences scoring as high). **(F)** *Microviridae* sequences clustered by their nearest one-hot encoded start/stop codon vectors relative to the ΦX174 reference. ΦX174 variants form distinct clusters with both high architecture similarity scores and high spike protein sequence identity to the ΦX174 spike protein.

**Figure S6.**
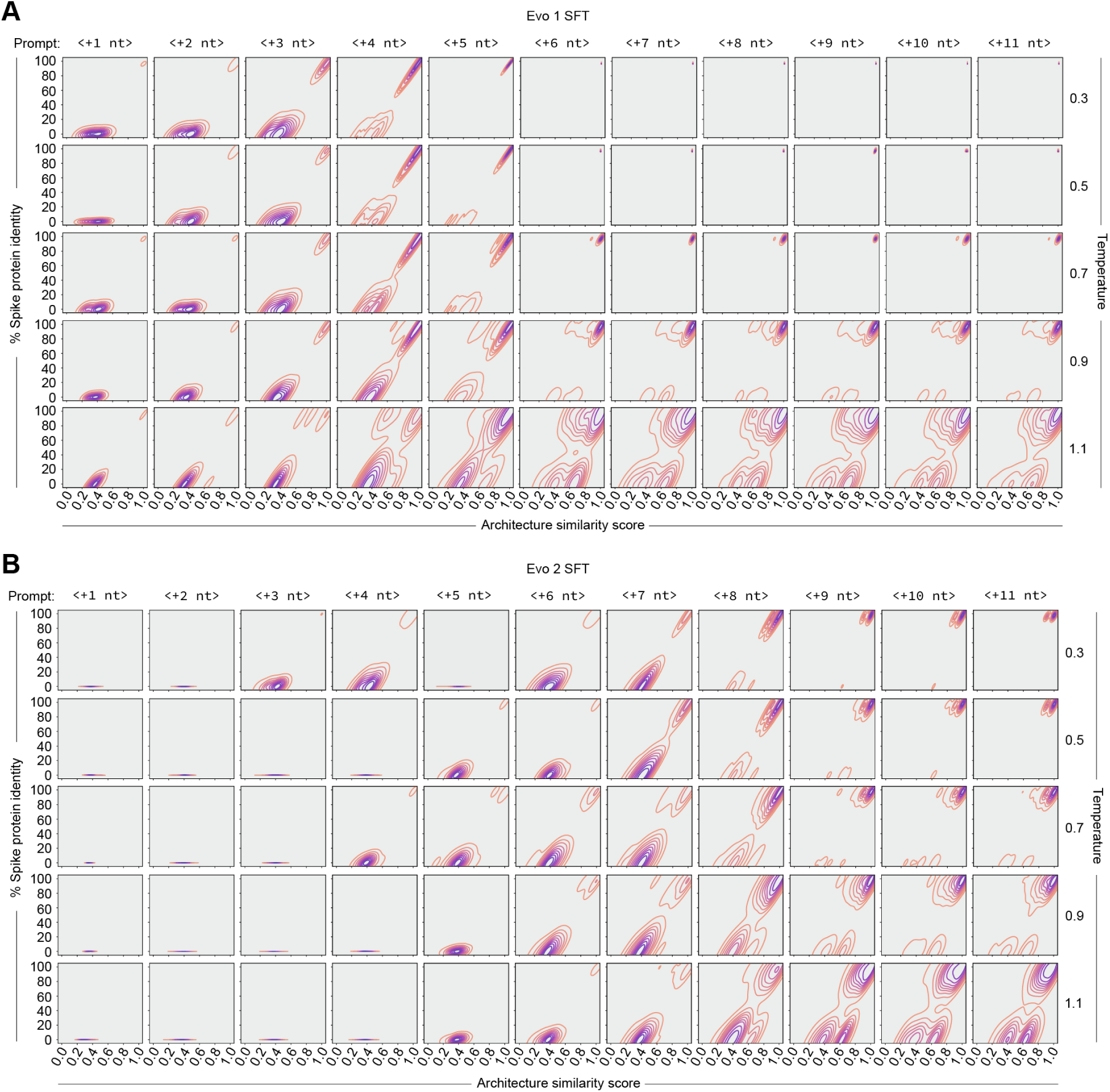
Generation temperature and prompt sweep. **(A)** Percent predicted spike protein identity and architecture similarity score of Evo 1 SFT- and Evo 2 SFT-generated sequences with increasing prompt lengths and generation temperatures. 𝑛 = 1000 sequences per parameter combination.

**Figure S7.**
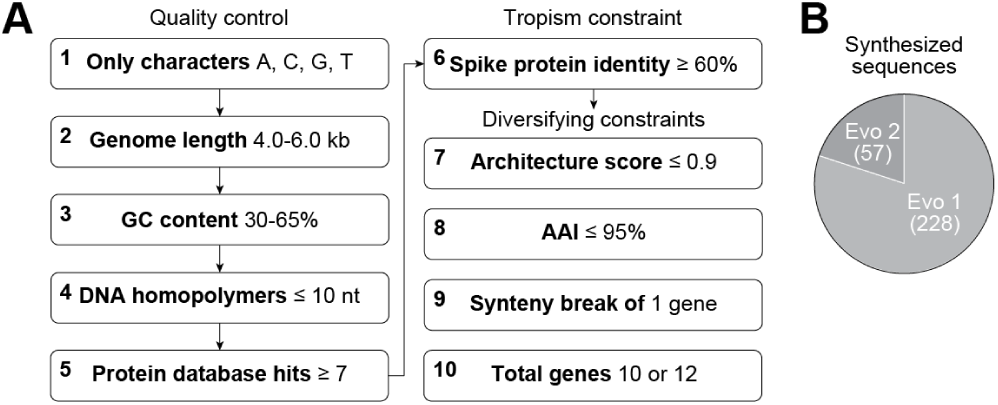
Supplementary data for bacteriophage genome design filtering. **(A)** Final filtering criteria for generated phage genomes. Diversifying constraints were preferred but not always applied. kb, kilobase; nt, nucleotides; AAI, average amino acid sequence identity. **(B)** Proportions of final sequence candidates generated by Evo 1 and Evo 2 that could be synthesized.

**Figure S8.**
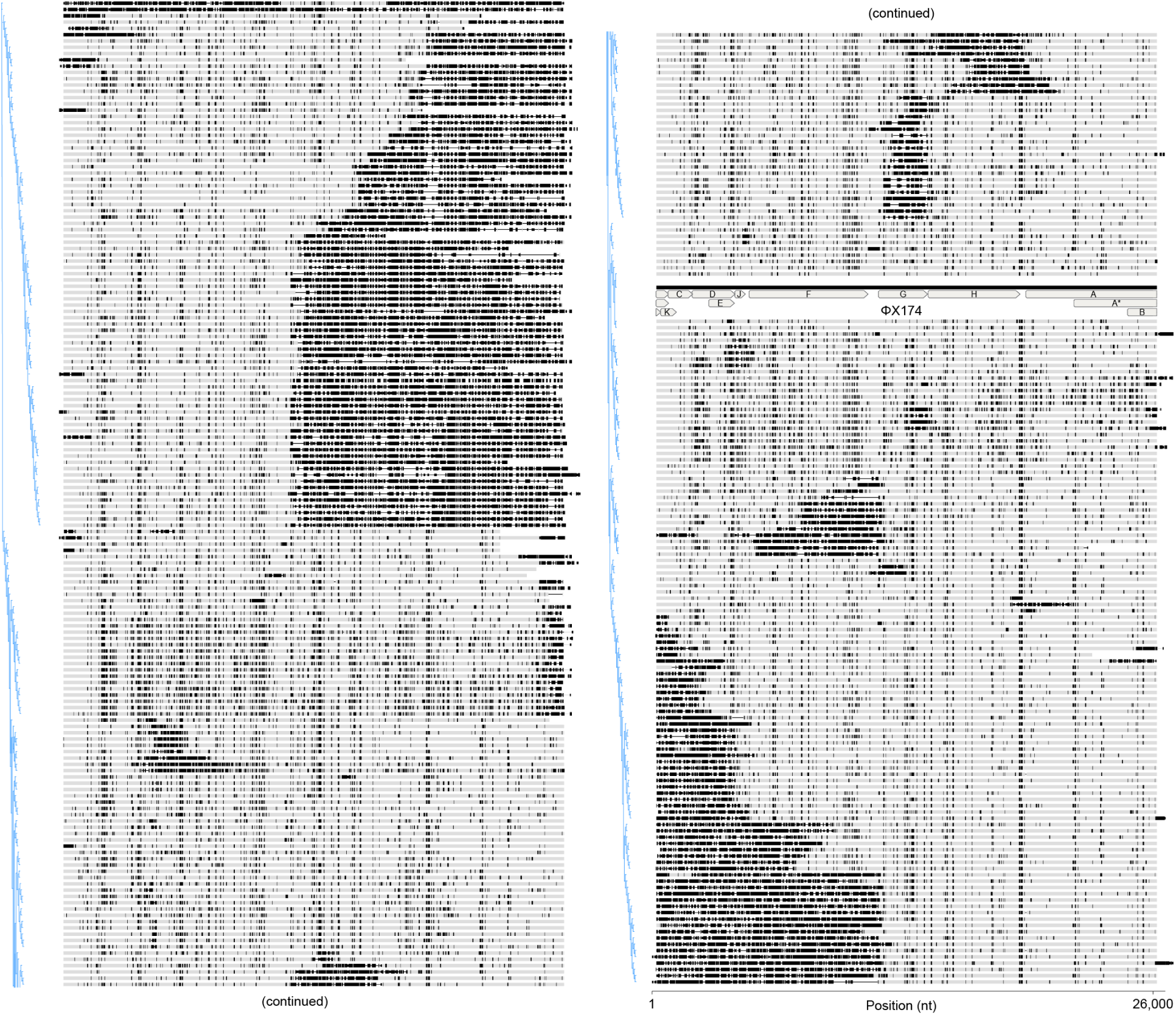
Alignment of synthesized bacteriophage genome design candidates. Alignment of final bacteriophage genome candidates and ΦX174, ordered by a neighbor-joining phylogenetic tree. Black, unaligned nucleotides to ΦX174, gray, aligned nucleotides to ΦX174.

**Figure S9.**
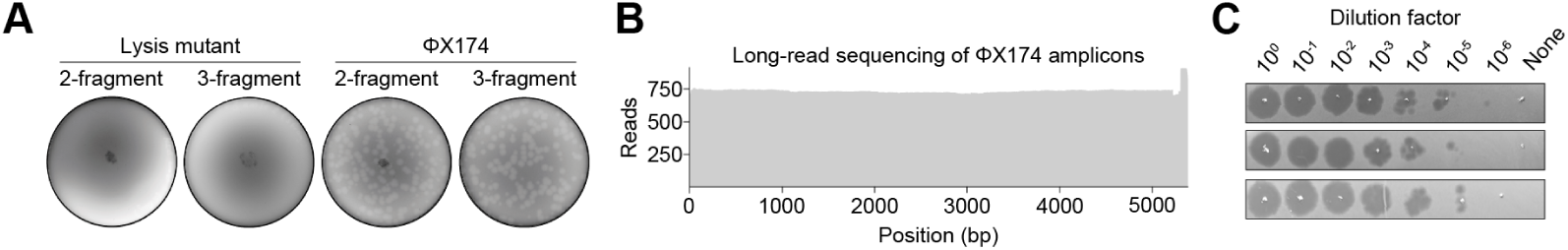
Supplementary data for bacteriophage genome assembly and rebooting. **(A)** Plaque assays of ΦX174 wild-type or lysis mutant genome assemblies transformed into *E. coli* C show robust phage rebooting with wild-type only. Two- and three-fragment Gibson assemblies were tested. **(B)** Long-read sequencing read counts of PCR amplicons from ΦX174 plaques. **(C)** Titrations of purified ΦX174 particles on *E. coli* C.

**Figure S10.**
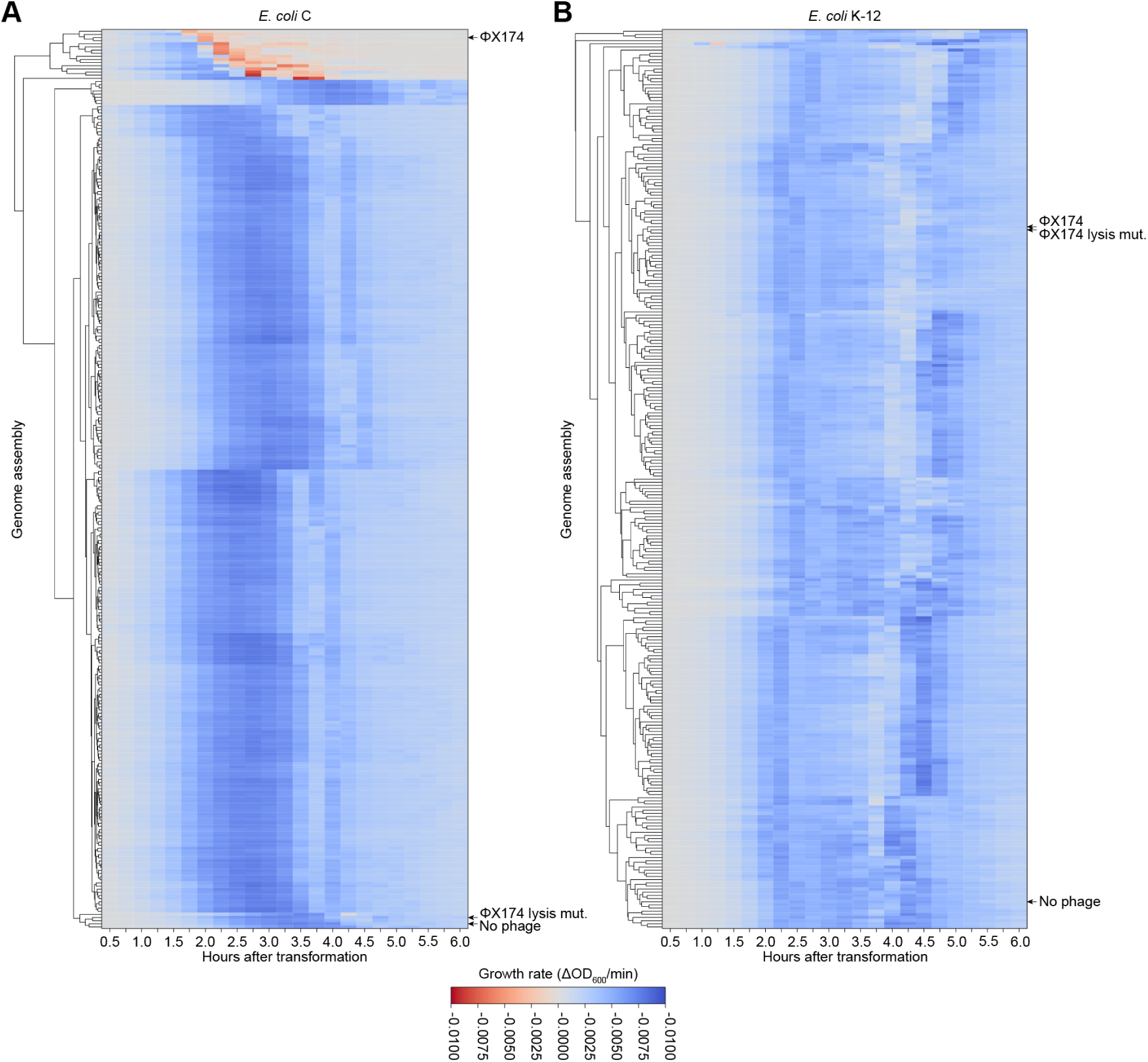
Supplementary data for bacteriophage rebooting in *E. coli* C and *E. coli* K-12. (A–B) Heatmaps of growth rate of *E. coli* C **(A)** and *E. coli* K-12 **(B)** over 6 hours after transformation with assembled phage genomes. Growth rates of *E. coli* C transformed with viable generated phages clusters with ΦX174.

**Figure S11.**
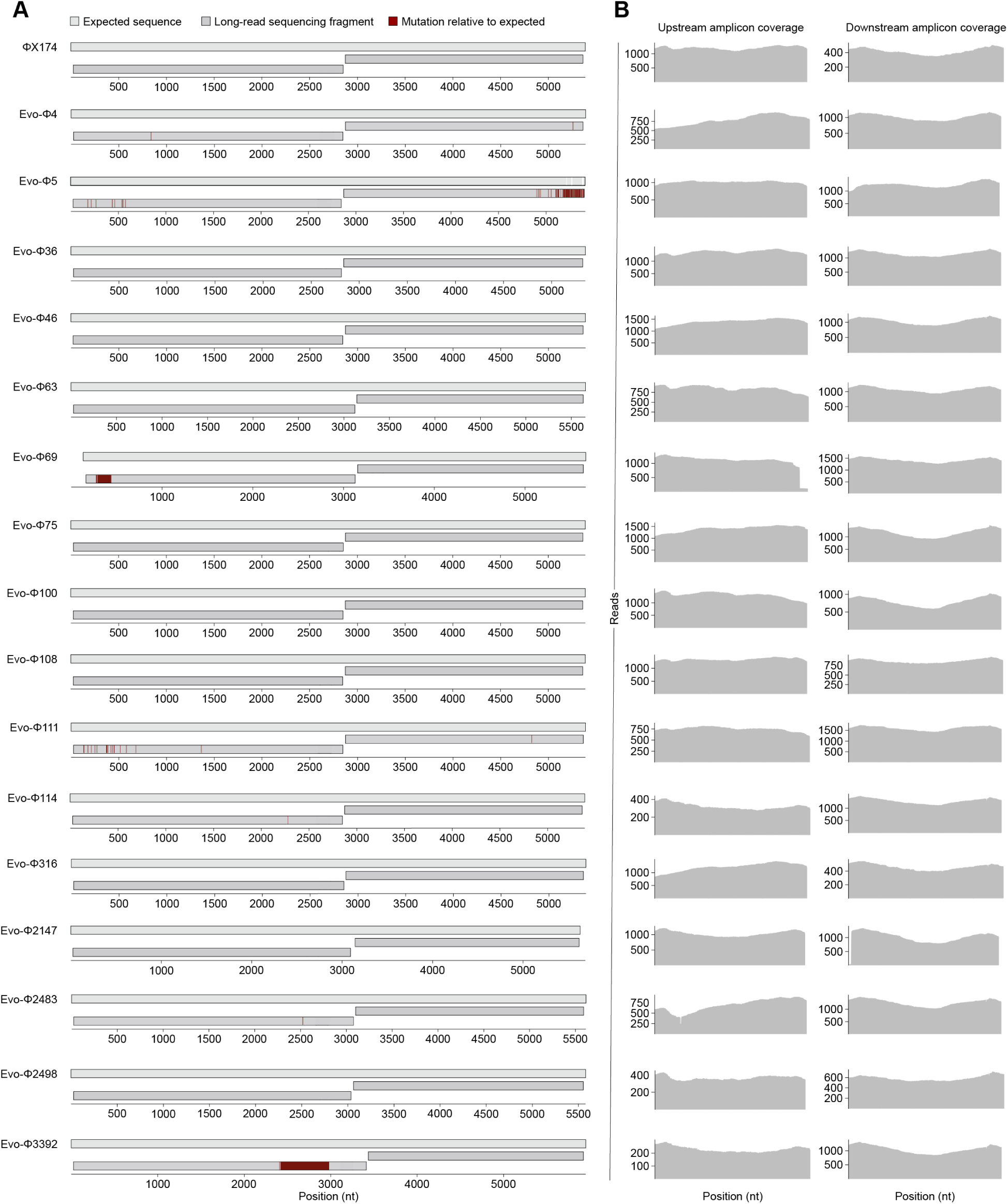
Long-read sequencing of viable generated bacteriophage genomes. **(A)** Long-read sequencing of PCR amplicons (dark gray) from ΦX174 and generated phage stocks, aligned against the expected sequence (light gray). Red, mutations relative to the expected sequence; white, gaps in the alignment. **(B)** Read counts from long-read sequencing.

**Figure S12.**
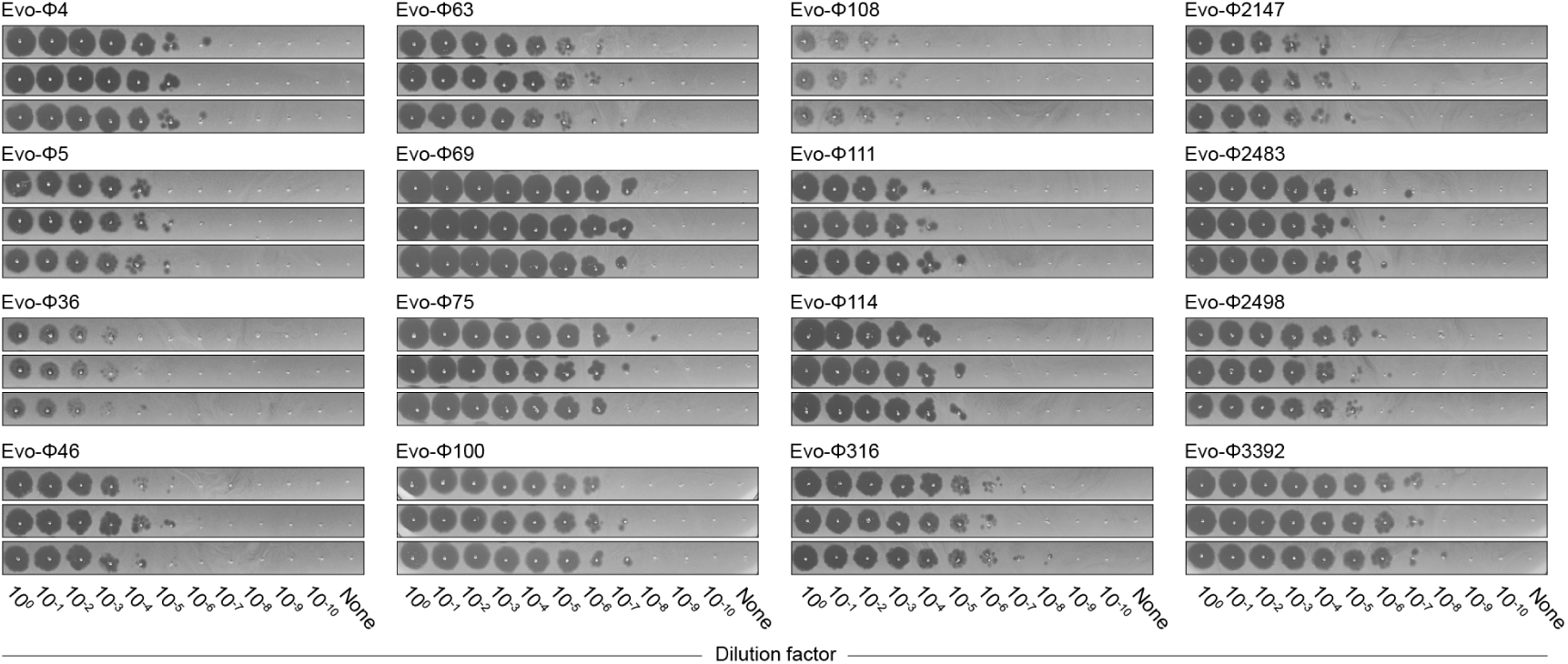
Titration of viable generated bacteriophage genomes. Plaque titrations of purified generated phages on *E. coli* C. Each row is a titration replicate.

**Figure S13.**
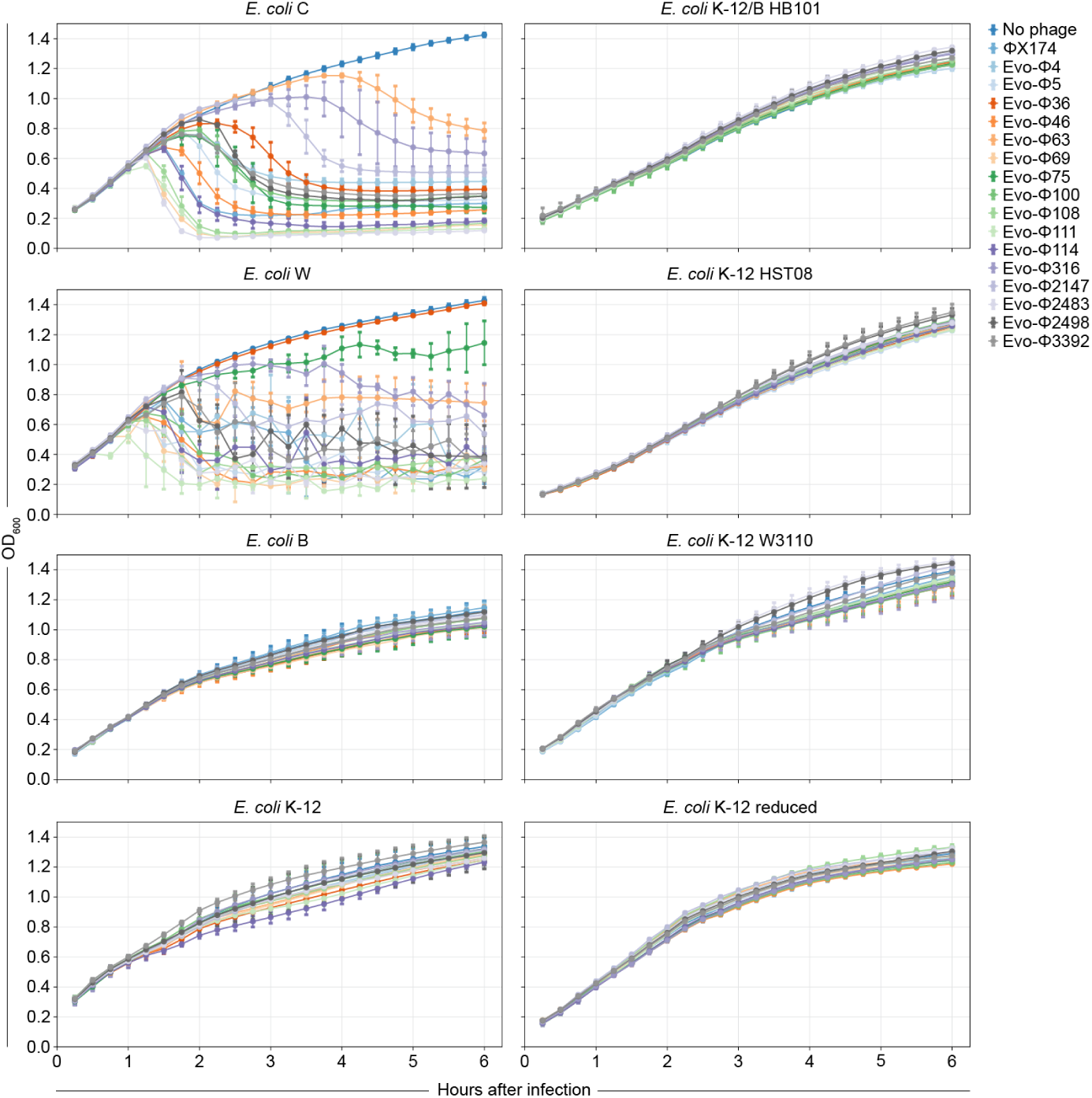
Supplementary data for host tropism assay. Growth curves of eight different *E. coli* strains infected with generated phages and ΦX174.

**Figure S14.**
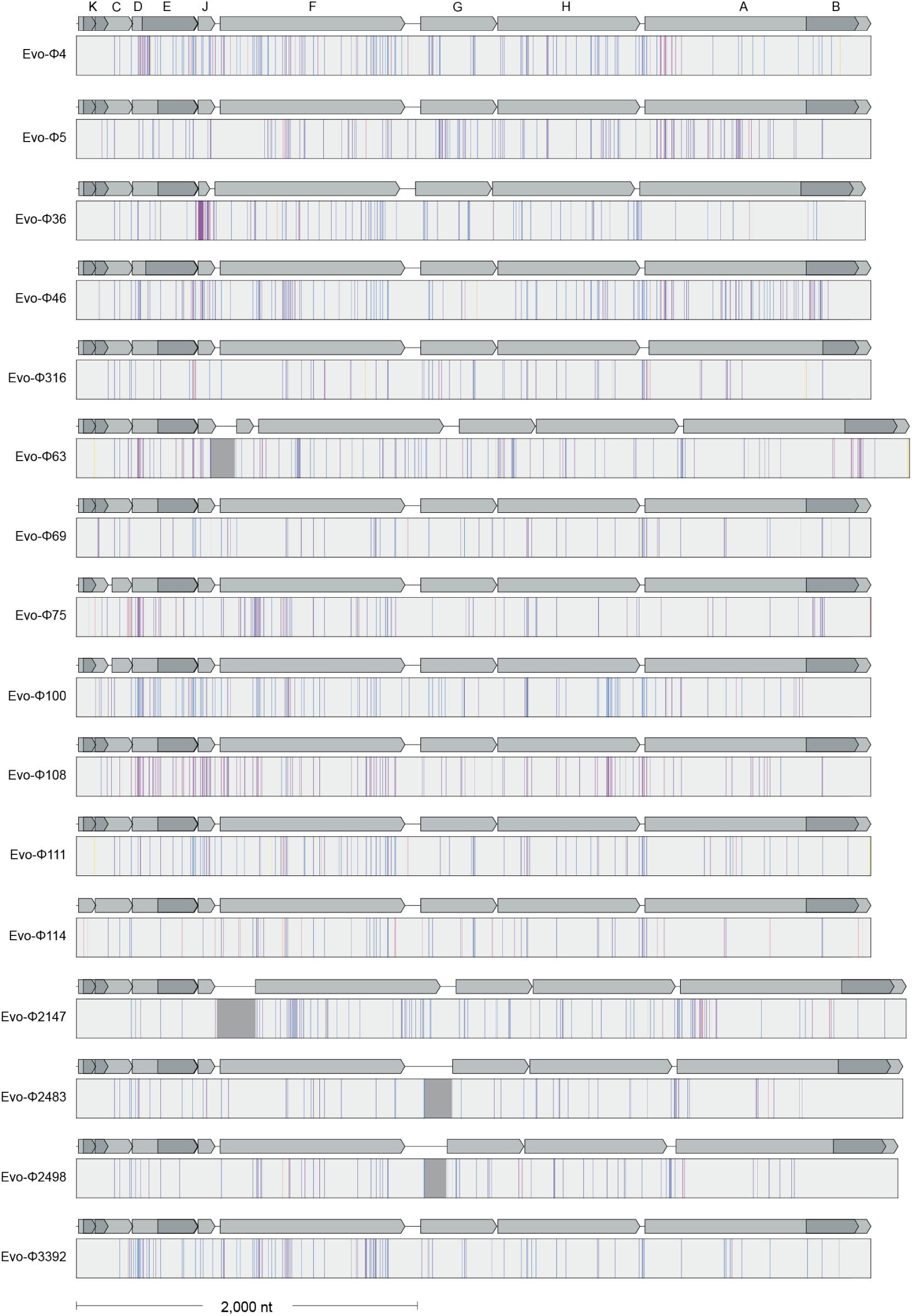
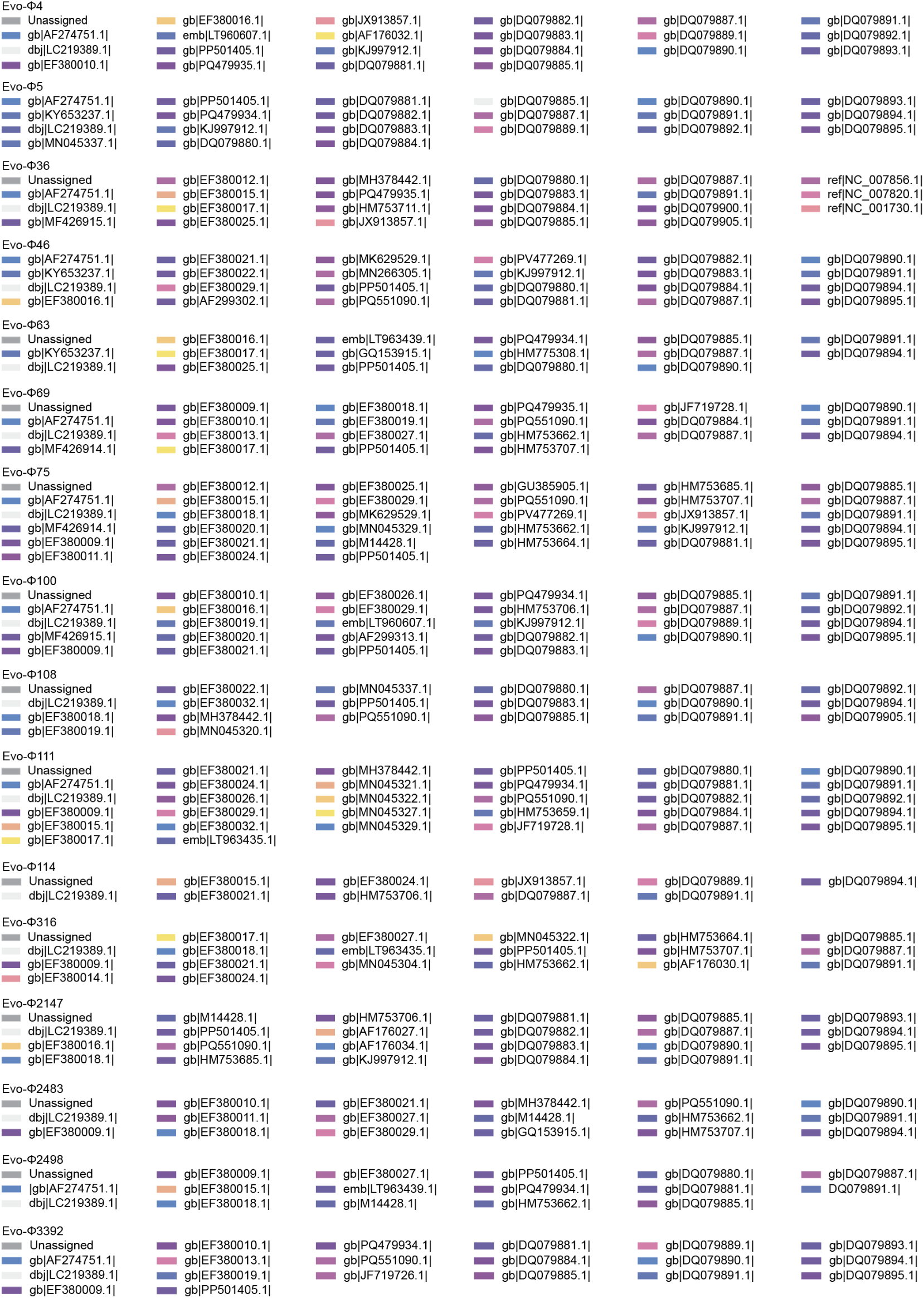
Supplementary data for cumulative genome attribution analysis. Sequence attribution analysis of generated phages, with nucleotides colored as “Unassigned” or by their top nucleotide BLAST hit in the core_nt database. Mutations in most generated phages cannot be completely attributed to mutations seen in nature.

**Figure S15.**
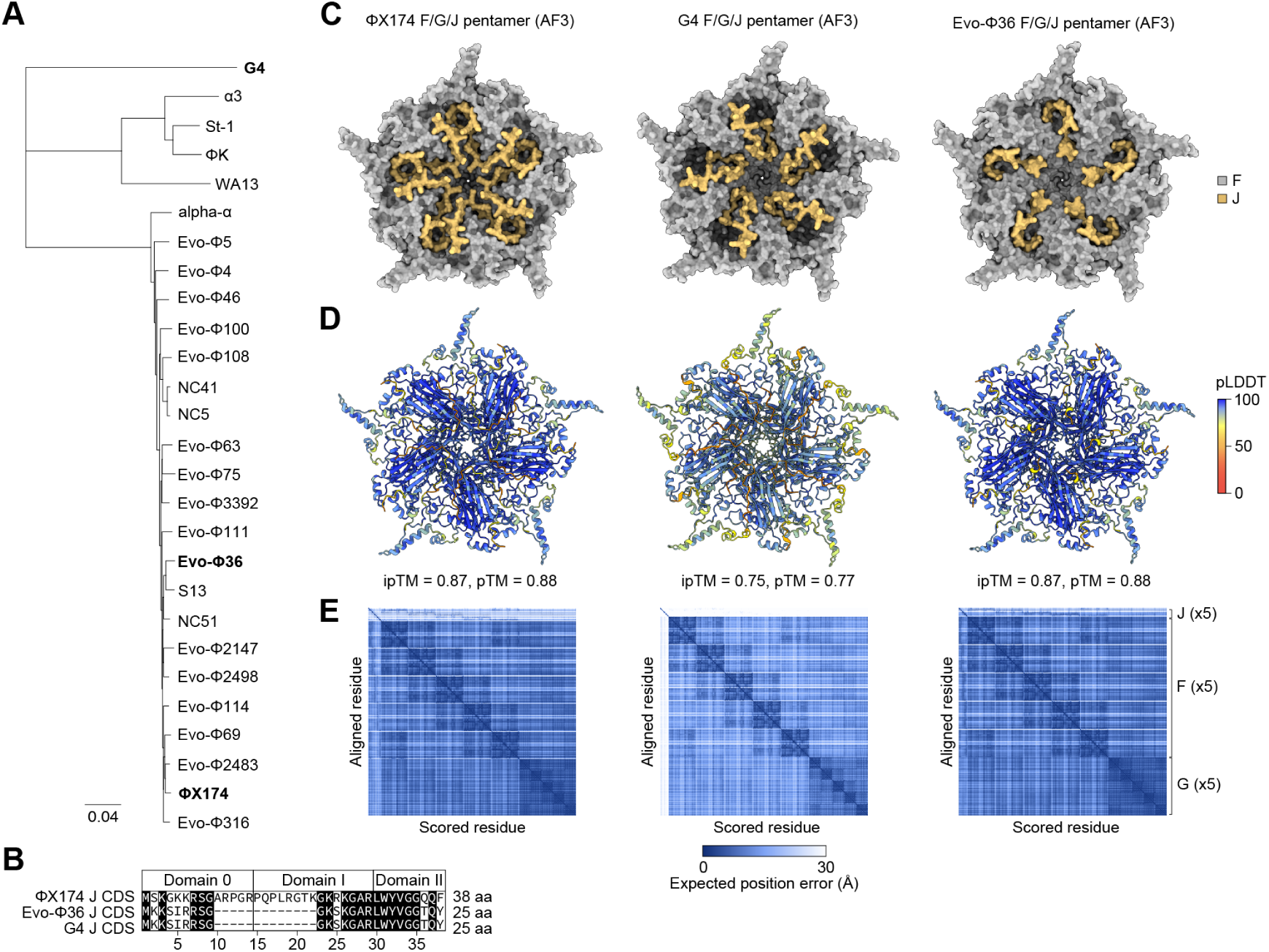
**Phylogenetic analysis of Evo-**Φ**36 J gene. (A)** Neighbor-joining phylogenetic tree of representative *Microviridae* phages and generated phages. **(B)** Coding sequence (CDS) of gene J of ΦX174, Evo-Φ36, and phage G4, with positions of domains 0, I, and II shown. Black shading, identical residues; aa, amino acids. **(C)** AlphaFold 3 (AF3) predictions of F (capsid, gray), spike (G, not visible), and J protein (yellow) pentamers of ΦX174, G4, and Evo-Φ36. The interior face of the pentamers is shown, highlighting the notable differences in J protein interactions with capsid proteins. **(D)** AF3 predictions of F/G/J pentamers colored by predicted local distance difference test (pLDDT) score. ipTM, interface predicted template modeling score; pTM, predicted template modeling score. **(E)** Predicted aligned error (PAE) plots of the AF3 predictions.

**Figure S16.**
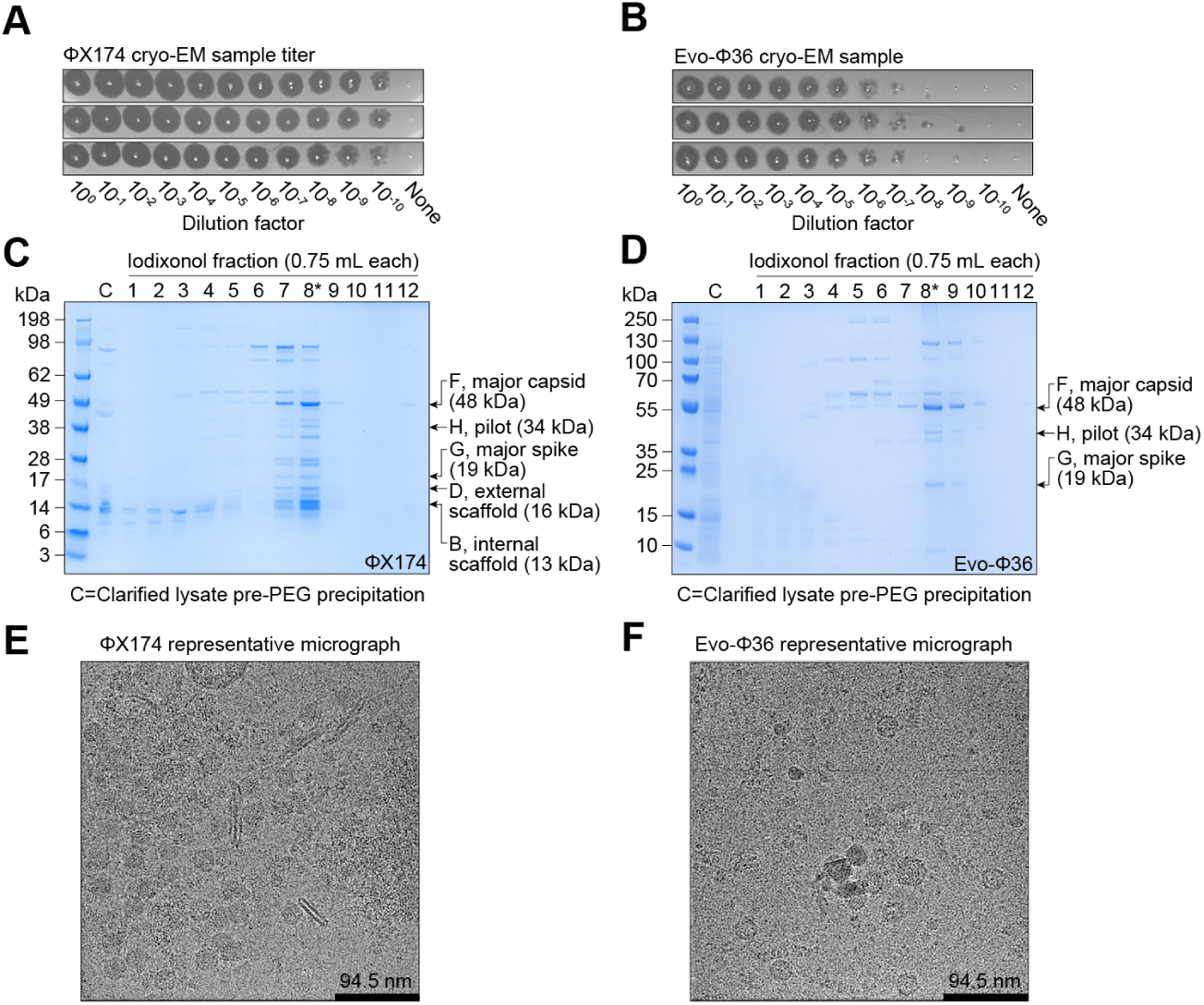
Supplementary data for cryo-EM analysis. (A–B) Titrations of purified ΦX174 **(A)** and Evo-Φ36 **(B)** cryo-EM samples. **(C–D)** SDS-PAGE of ΦX174 **(C)** and Evo-Φ36 **(D)** iodixonol gradient fractions. *, fraction used for cryo-EM. **(E–F)** Representative cryo-EM micrographs of ΦX174 **(E)** and Evo-Φ36 **(F)** grids used for cryo-EM data collection.

**Figure S17.**
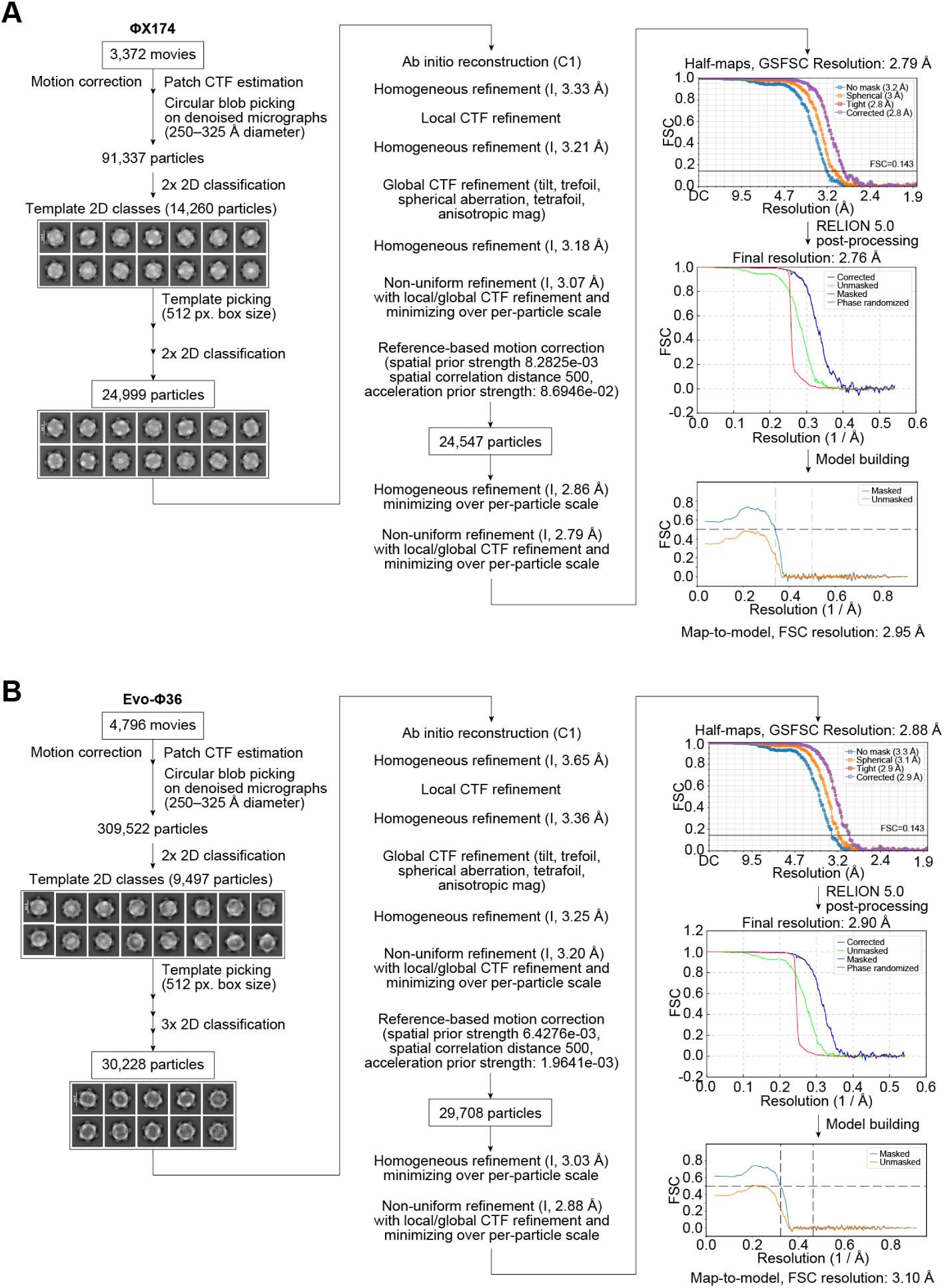
Cryo-EM data processing workflow. (A–B) Cryo-EM data processing workflows in cryoSPARC for ΦX174 **(A)** and Evo-Φ36 **(B)** with half-map FSCs as reported by cryoSPARC without auto-tightening of the mask, FSCs from RELION with threshold-based mask, and map-to-model FSCs from Phenix with a threshold of FSC = 0.5. Masks for RELION post-processing were constructed using a low-pass filter of 20 Å, 1 pixel extension, and 8 pixel soft-edge with a binarization threshold of 0.015 and 0.01 for ΦX174 and Evo-Φ36, respectively.

**Figure S18.**
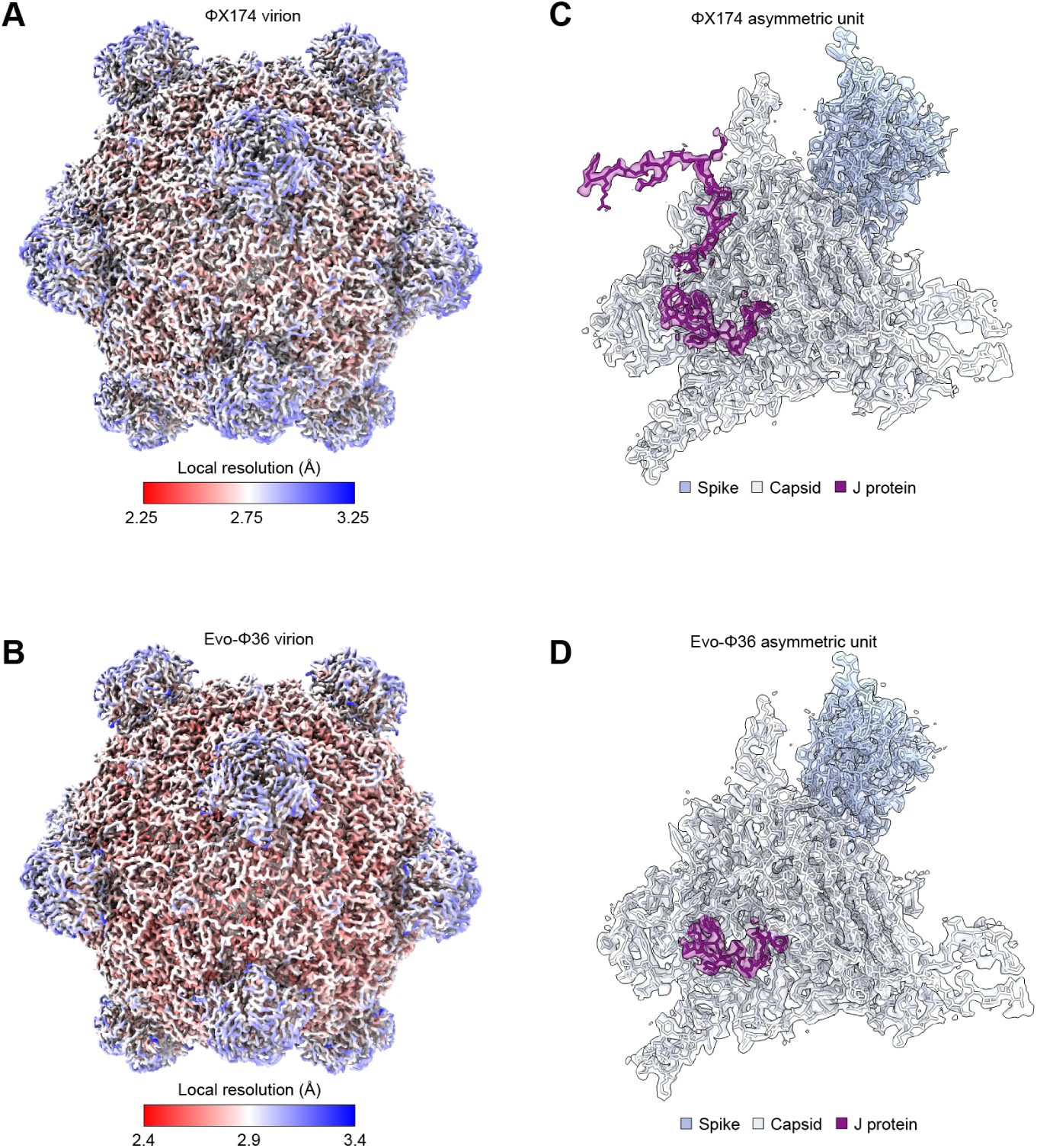
Additional validation of cryo-EM maps and atomic models. (A–B) Local resolution estimation for for ΦX174 **(A)** and Evo-Φ36 **(B)** as output by cryoSPARC (FSC = 0.5). **(C–D)** Visualization of atomic models fit into cryo-EM densities for an asymmetric unit of ΦX174 **(C)** and Evo-Φ36 **(D)**.

**Figure S19.**
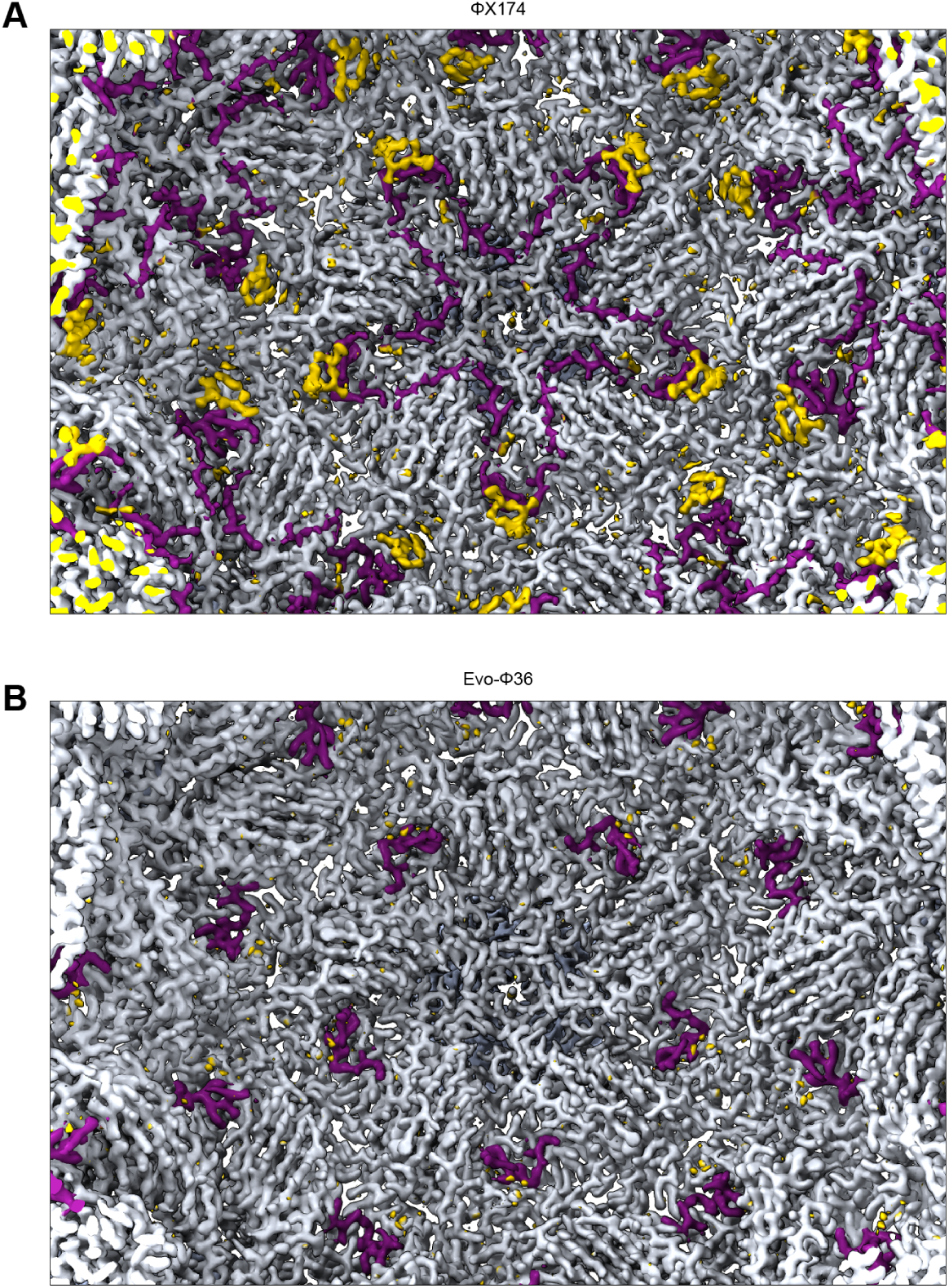
Interior surface views of cryo-EM density maps. (A–B) Interior surface view of the cryo-EM density maps of ΦX174 **(A)** and Evo-Φ36 **(B)**, colored by zone based on the model with capsid (F) in gray, spike (G) in light blue (largely not visible), and DNA packaging protein (J) in purple. Compare to Figure 4J. Gold corresponds to unmodeled densities, primarily of nucleic acids.

**Figure S20.**
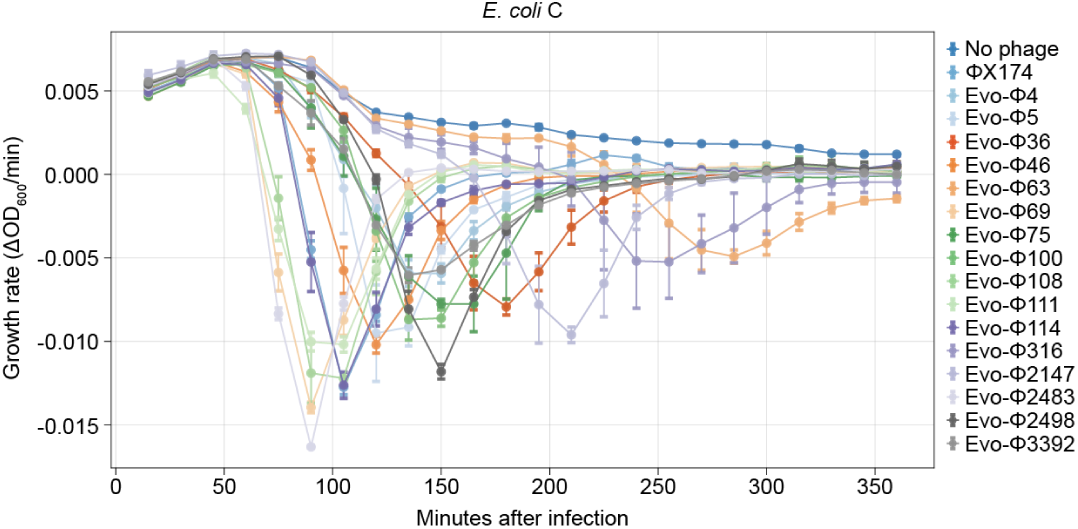
Supplementary data for lytic assay. Growth rates of *E. coli* C infected by generated phages and ΦX174, with no phage infection control. Data point, mean OD_600_ value; error bar, standard deviation; 𝑛 = 3 infection replicates.

**Figure S21.**
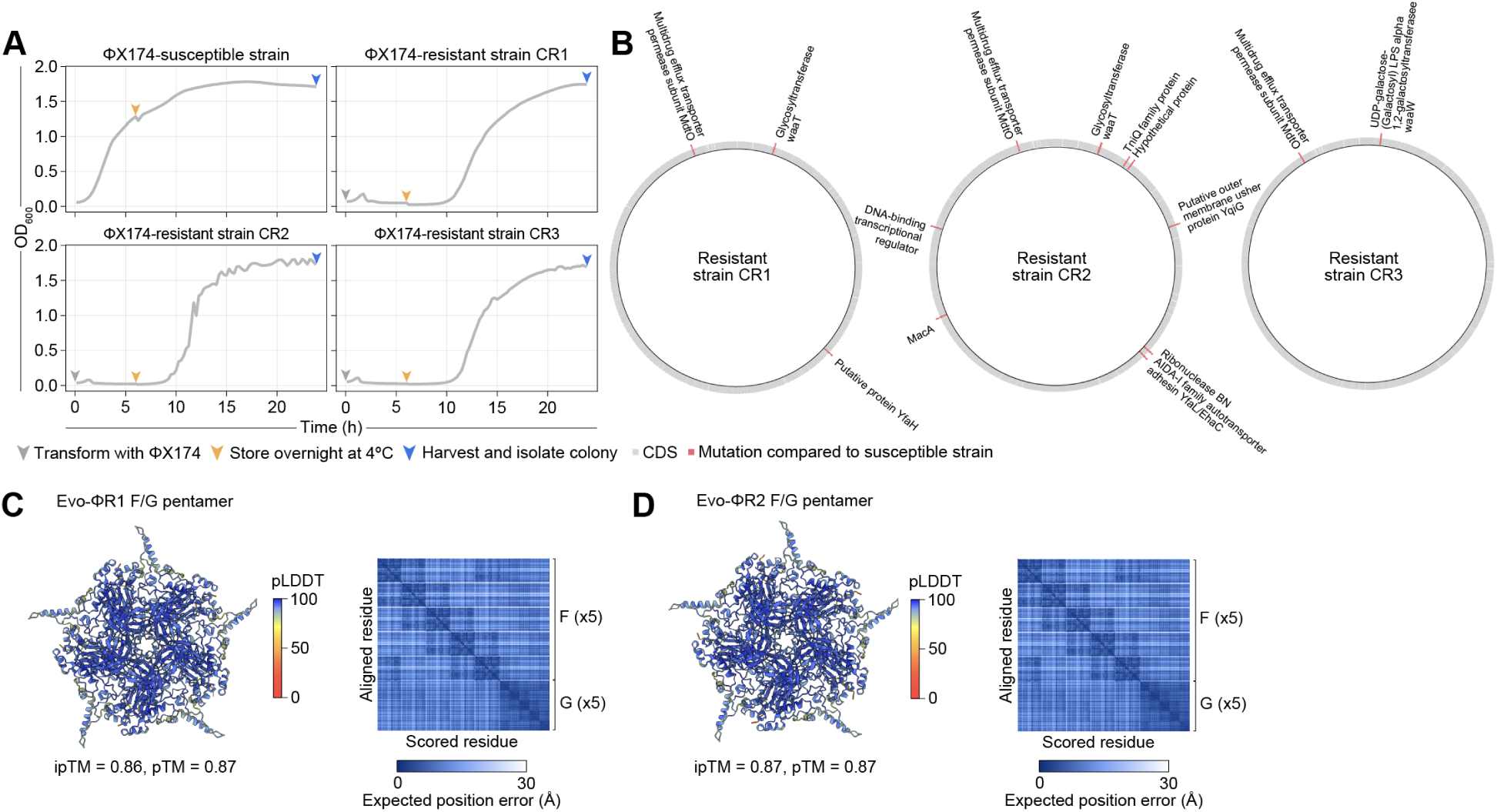
Supplementary data for bacteriophage resistance assay. **(A)** Growth curves of *E. coli* C without infection by ΦX174 (ΦX174-susceptible strain), or transformed with ΦX174 genome (ΦX174-resistant strain). Arrowheads indicate key timepoints for producing and harvesting the different strains. **(B)** Whole-genome sequencing of resistant strains CR1, CR2, and CR3 show unique mutations compared to the susceptible strain. Gray, predicted coding sequence (CDS); red, point mutation relative to susceptible strain. **(C–D)** AlphaFold 3 (AF3) predictions of F (capsid) and spike (G) pentamers of Evo-ΦR1 and Evo-ΦR2. The exterior face of the pentamers is shown, colored by predicted local distance difference test (pLDDT) score. Predicted aligned error (PAE) plots of the AF3 predictions are shown on the right. ipTM, interface predicted template modeling score; pTM, predicted template modeling score.

**Table S1.**
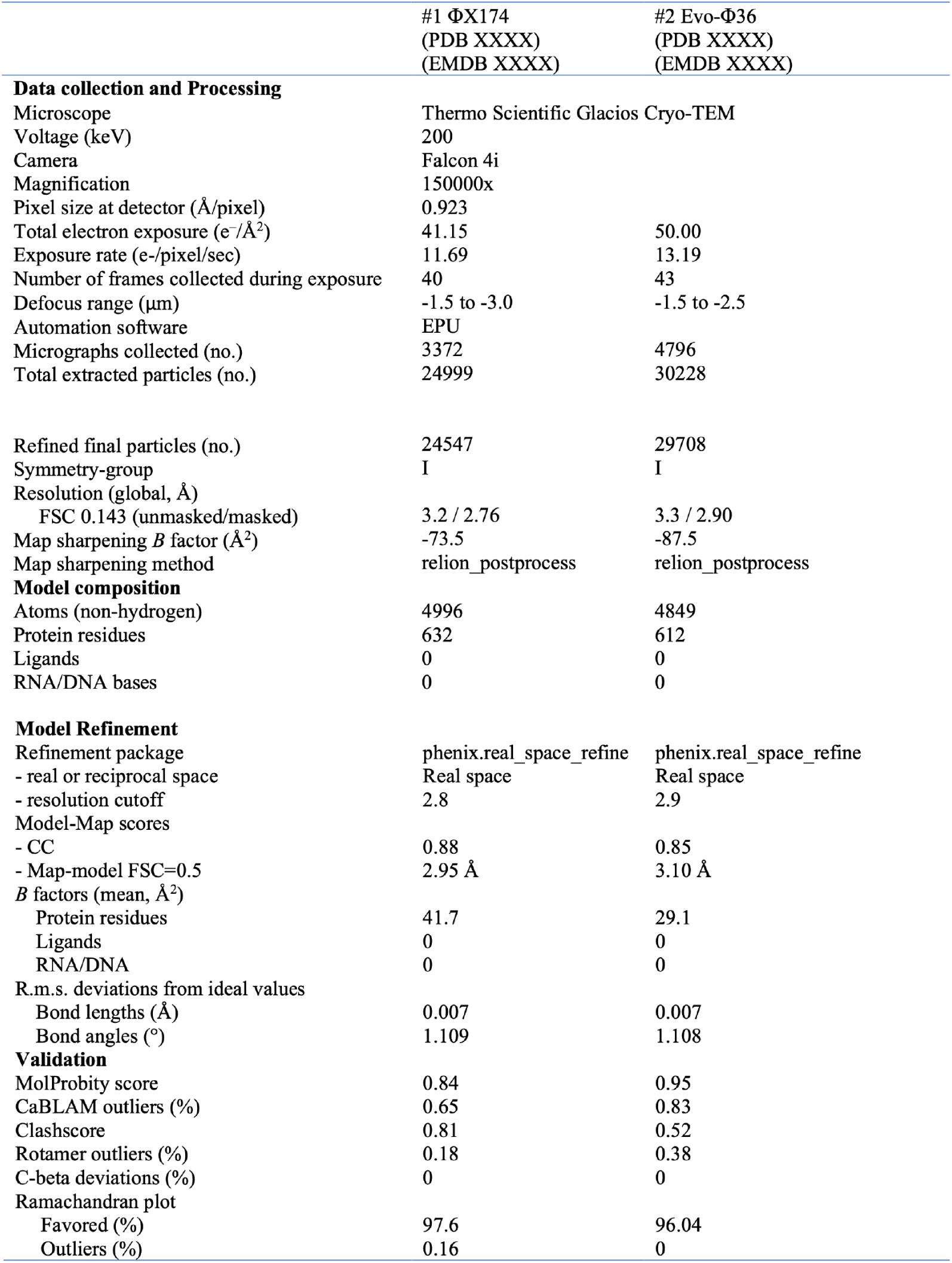
Cryo-EM data collection, refinement, and validation statistics.

## **D.** Supplementary files

### **D.1.** File S1

Generated bacteriophage genome candidates, metrics, and assembly fragments DNA oligo sequences Bacteriophage genome amplicons for long-read sequencing Functional sequencing-verified bacteriophage genome sequences

